# A dynamin-like protein in the human malaria parasite plays an essential role in mitochondrial homeostasis and fission during asexual blood stages

**DOI:** 10.1101/2024.03.25.586598

**Authors:** Vandana Thakur, Shweta Singh, Sumit Rathore, Azhar Muneer, Md. Muzahidul Islam, Gaurav Dutta, Mudassir M. Banday, Priya Arora, Mohammad E. Hossain, Shaifali Jain, Shakir Ali, Asif Mohmmed

## Abstract

Malaria parasite harbors a single mitochondrion and its proper segregation during the parasite multiplication is crucial for propagation of the parasite within the host. Mitochondrial fission machinery consists of a number of proteins that associate with mitochondrial membrane during segregation. Here, we have identified a dynamin-like protein in *P. falciparum*, *Pf*Dyn2, and deciphered its role in mitochondrial division, segregation and homeostasis. GFP targeting approach combined with high resolution microscopy studies showed that the *Pf*Dyn2 associates with mitochondrial membrane to form a clip/hairpin loop like structure around it at specific sites during mitochondrial division. The C-terminal degradation tag mediated inducible knock-down (iKD) of *Pf*Dyn2 resulted in significant inhibition of parasite growth. *Pf*Dyn2-iKD hindered mitochondrial development and functioning, decreased mtDNA replication, and induced mitochondrial oxidative-stress, ultimately causing parasite death. Further, treatment of parasites with dynamin specific inhibitors disrupted the recruitment of *Pf*Dyn2 on the mitochondria, blocked mitochondrial development, and induced oxidative stress. Regulated overexpression of a phosphorylation mutant of *Pf*Dyn2 (Ser-612-Ala) had no effect on the recruitment of *Pf*Dyn2 on the mitochondria; normal mitochondrial division and parasite growth showed that phosphorylation/dephosphorylation of this conserved serine residue (Ser612) may not be responsible for regulating recruitment of *Pf*Dyn2 to the mitochondrion. Overall, we show essential role of *Pf*Dyn2 in mitochondrial dynamics and fission as well as in maintaining its homeostasis during asexual cycle of the parasite.

## Introduction

In spite of remarkable developments in human healthcare system, malaria remains to be a global burden mainly in the tropical and subtropical regions causing substantial morbidity and mortality. Globally, malaria resulted in >240 million cases leading to ∼ 619,000 mortalities in the year 2021, majority of which these deaths are among children aged below 5 years [1]. Among five different *Plasmodium* parasite, *P. falciparum* causes most fatal form of disease, the cerebral malaria. Considerable improvement has been made to control malaria in the last decade, due to availability and effectiveness of artemisinin combination therapy (ACT) [2][3]. Unfortunately, in the wake of emergence of parasite lines resistant to commonly used anti-malarial, including artemisinin, there is urgent need to identify new drug-targets and develop novel anti-malarial drugs.

Parasite harbours two organelles of prokaryotic origin, the relict plastid apicoplast and the mitochondrion, which cannot be formed de novo by the parasite, therefore, all daughter merozoites carry a copy of these organelles. Mitochondria play role in producing energy (ATP) and also play diverse roles in various cellular activities, regulating a multitude of cellular functions, which include Ca2+ homeostasis, cellular signaling, reactive oxygen species (ROS) production, and apoptotic cell-death innate immunity, autophagy and stem cells reprogramming [4], [5], [6]. Mitochondrial vary in shape, size and morphology from small granules to tubular structure in an organized fashion to maintain fusion and fission events [7], [8]. Mitochondrial division and segregation during cell cycle, including fission and fusion events, is regulated by several GTPases: dynamin-related proteins (DRPs)[9]. Mitochondrial fission is the vital process for the cell survival, as it segregate daughter mitochondria through cell division, calcium signaling, allocate mtDNA, segregate damaged mitochondria for degradation (mitophagy) and initiate apoptosis when necessary [10], [11]. In eukaryotes, Drp1 is the key player in the fission machinery which exists as dimers or tetramers in cytoplasm[12]. This unique mechanochemical enzyme, characterized by a large GTPase domain, gets typically recruited to the outer membrane of mitochondria through its interaction with other components of the mitochondrial fission machinery including Fis1 and Mdv1 [13][14]. After its recruitment, Drp1 gets assembled into higher-order complexes (oligomers) at endoplasmic reticulum (ER)/actin-mitochondrial contact sites and forms clip-like structure around the mitochondria to induce mitochondrial fission through its GTPase activity [15][16]. Interestingly, ER-mitochondrial interactions are important not only for the regulation of mitochondrial morphology, but also for the regulation of intracellular Ca^2+^ signals [22]. The imbalance in fission machinery results in highly interconnected mitochondrial instead of tubular structures in mammalian and yeast cells [17], [18], [19]. Under these conditions, mitochondria were shown to produce excessive amounts of reactive oxygen species (ROS) that can disrupt the organelle function in terms of ATP synthesis [20], [21]. Increased levels of mitochondrial ROS can promote excessive Ca2+ uptake, oxidative imbalance and protein misfolding, leading to a loss in membrane potential and subsequently causing the release of pro-apoptotic factors that can initiate cell death signaling [22][20].

The role of Drp1 is mediated by several post-translational modifications, such as ubiquitylation, SUMOylation, S-nitrosylation, and phosphorylation [9]. Drp1 phosphorylation is the most studied post-translation modification that regulating mitochondrial fission, and it occurs at two different serine residues: S616 and S637 (in human Drp1 isoform 1) which are regulatory for mitochondrial fission and fusion in the cell respectively [23]. During the cell cycle Drp1-S616 phosphorylation via cdk1/cyclin B kinase induce mitochondrial fission [24]. whereas phosphorylation of S637, via cAMP-dependent protein kinase A (PKA), inhibits mitochondrial fission. During nutrient deprivation/cell-death phosphorylation of S637 protect mitochondria from autophagosomal degradation, by impairing Drp1 GTPase activity and preventing translocation of Drp1 to mitochondria [4]. Conversely, dephosphorylation of this residue is carried out by the calcium-dependent phosphatase calcineurin, which increases mitochondrial recruitment of Drp1, in turn promoting mitochondrial fission leading to cell death and necrosis [23][9].

*Plasmodium* harbours single mitochondrion in each unicellular parasite; It has been shown to associate with ER and apicoplast in the parasite [25], [26], [27]During the parasite division cycle, fission and segregation of mitochondria are well coordinated with the nuclear division, segregation of other organelles, and cytokinesis [28][29]. However, very little is known about the machinery involved in mitochondrial fission in the malaria parasite. The genome of the *P. falciparum* parasite encodes two dynamin-like proteins, dynamin-like protein-1 (*Pf*Dyn1 Gene ID: PF3D7_1145400) and dynamin-like protein-2 (*Pf*Dyn2 Gene ID: PF3D7_1037500); in addition, there is another putative dynamin protein (PF3D7_1218500), however, it does not harbour all the domains required to be a dynamin protein. Previous work suggests that *Pf*Dyn1 might be involved in the process of nutrient ingestion or endocytosis [30], [31]. Based upon molecular architecture and phylogenetic analysis, *Pf*Dyn2 was placed in the dynamin like proteins (Dlps) subgroup that might be involved in vesicular trafficking and/or organelle segregation [32], [33]. However, the exact role of *Pf*Dyn2 has yet not been characterised in *P. falciparum*. In the present study, we have functionally characterized homolog of dynamin in *P. falciparum Pf*Dyn2 and deciphered its essentiality in mitochondrial dynamics and parasite survival. Understanding of unique metabolic pathways, which are functionally vital for maintenance and homeostasis of mitochondrial organelles, may help us to design novel antimalarial strategies.

## Results

### Expression and subcellular localization of *Pf*DYN2-GFP fusion protein in transgenic parasites

The *Pf*Dyn2 is a 709 amino acids long protein that harbor three functional domains: Dynamin-N (29-209 aa; Pfam E-value 2.4e^-45^), Dynamin M Central domain (2210-508 aa; Pfam E-value 7.5 e^-85^) and GTPase effector Domain (GED) (615-704 aa; Pfam E-value 8.3e^-27^) (Fig 1A, S1). To understand putative role of *Pf*DYN2 in the parasite, we generated transgenic parasites expressing *Pf*Dyn2-GFP fusion protein. Expression of fusion protein was confirmed by western blot analysis of the total lysates of the transgenic parasites using anti-GFP antibodies; the anti-GFP antibody detected a band of ∼102 kDa, corresponding to the fusion protein, specifically in the transgenic parasites (Fig. 1B, S3). The transgenic parasites were used for fluorescence and confocal microscopy studies to localize the *Pf*Dyn2-GFP fusion protein in different developmental stages of the asexual cycle. In early stages, the GFP was observed to be in single foci in the cytosol of the parasite. In the mid trophozoite stages, the fluorescence was localized in a number of punctate structures in the parasite cytosol. In late-trophozoite stages, several fluorescent punctate like structures were observed distributed in the parasite cytosol (Figs. 1C, S2). In schizont stage, these fluorescence foci were found close to the nucleus.

**Figure 1:**
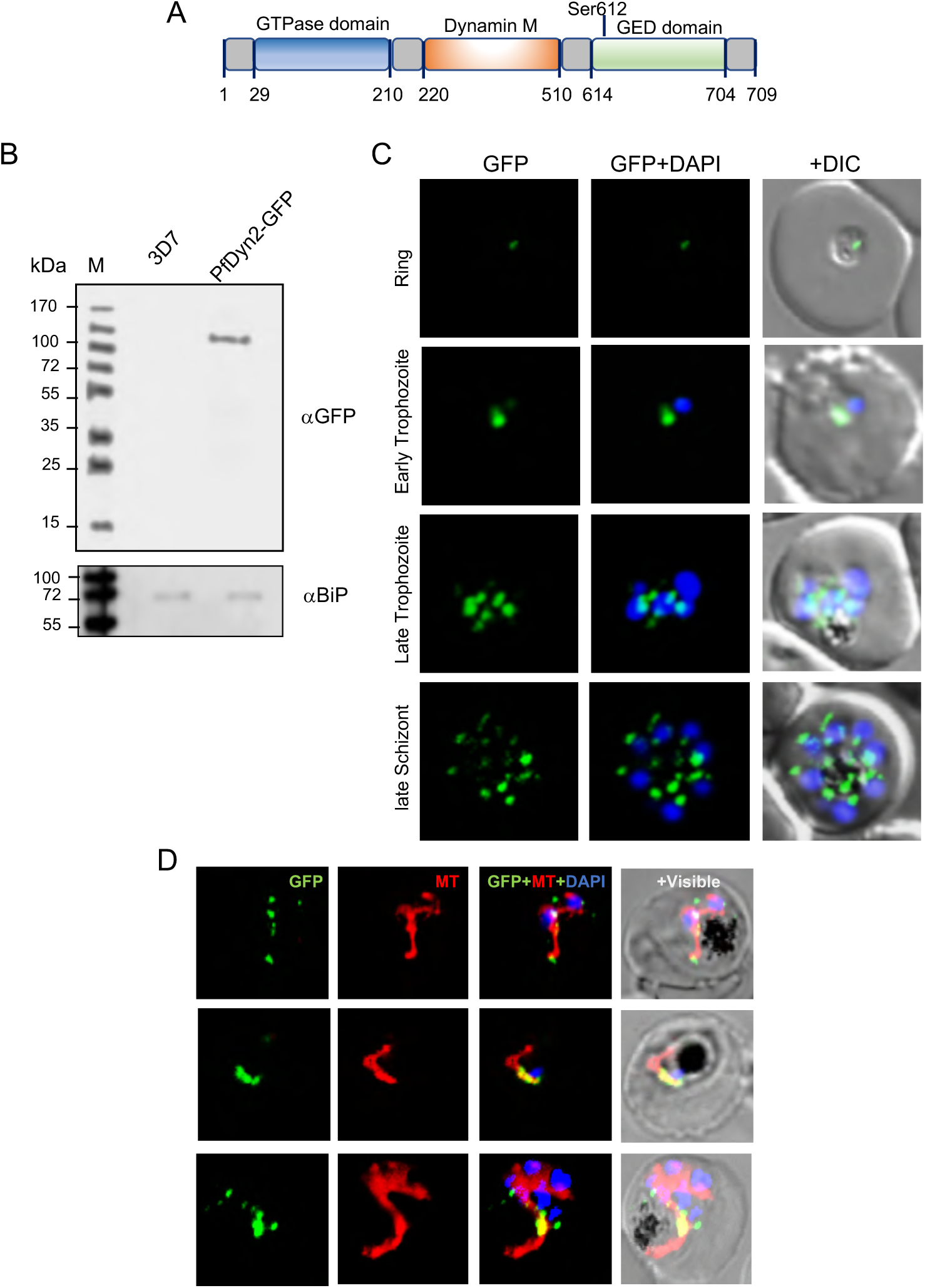
Expression and localization of *Pf*Dyn2-GFP fusion protein in transgenic parasites. (A) Schematic showing domain organization of *Pf*Dyn2 that harbours three functional domains: Dynamin-N (29-209 aa), Dynamin M Central domain (2210-508 aa), and GTPase effector Domain (GED) (615-704 aa). A conserved serine residue, associated with regulatory phosphorylation linked with mitochondrial fission, is marked. (B) Expression and localization of *Pf*Dyn2-GFP fusion protein through asexual parasite developmental stages (ring, trophozoite, and schizont stages); the parasite nuclei are stained by DAPI (blue) and parasites were visualized by a confocal laser scanning microscope. (C) Western blot analysis of lysate of transgenic and wild-type parasites using an anti-GFP antibody. The fusion protein band (∼102 kDa) was detected in the transgenic parasites only (lane 1) and not in the wild-type parasites (lane 2). Blot ran in parallel with an equal amount of the same sample, probed with anti-BiP antibody was used as a loading control. (D) Fluorescence image of *Pf*Dyn2-GFP parasites stained with MitoTracker (red). Parasites nuclei were stained with DAPI and the parasite was visualized by confocal microscope.

### *Pf*Dyn2 is membrane bound protein which associates with mitochondria

To further understand the localization of *Pf*DYN2 and its possible association with organelles in the parasite, detailed organelle labelling and localizations confocal studies were carried out. The parasites were stained with MitoTracker CMXRos (Red), a fluorescent dye that specifically labels the active mitochondria in live cells. In these parasites, partial co-localization was observed between GFP foci and mitochondria at the late stages, when mitochondria were about to divide (Fig. 1D, S3 and S4). To understand these localizations in detail, co-stained parasites were analyzed using Structured Illumination Microscopy (SIM); the SIM images showed that GFP foci at discrete locations overlapping with the mitochondria at the branching division sites (Fig. 2A, B); the 3D reconstruction of Z-stack images clearly showed that the *Pf*DYN2-GFP fusion protein associate with mitochondria surface and form clip-like structure at the time of mitochondrial division (Fig. 2C).

**Figure 2.**
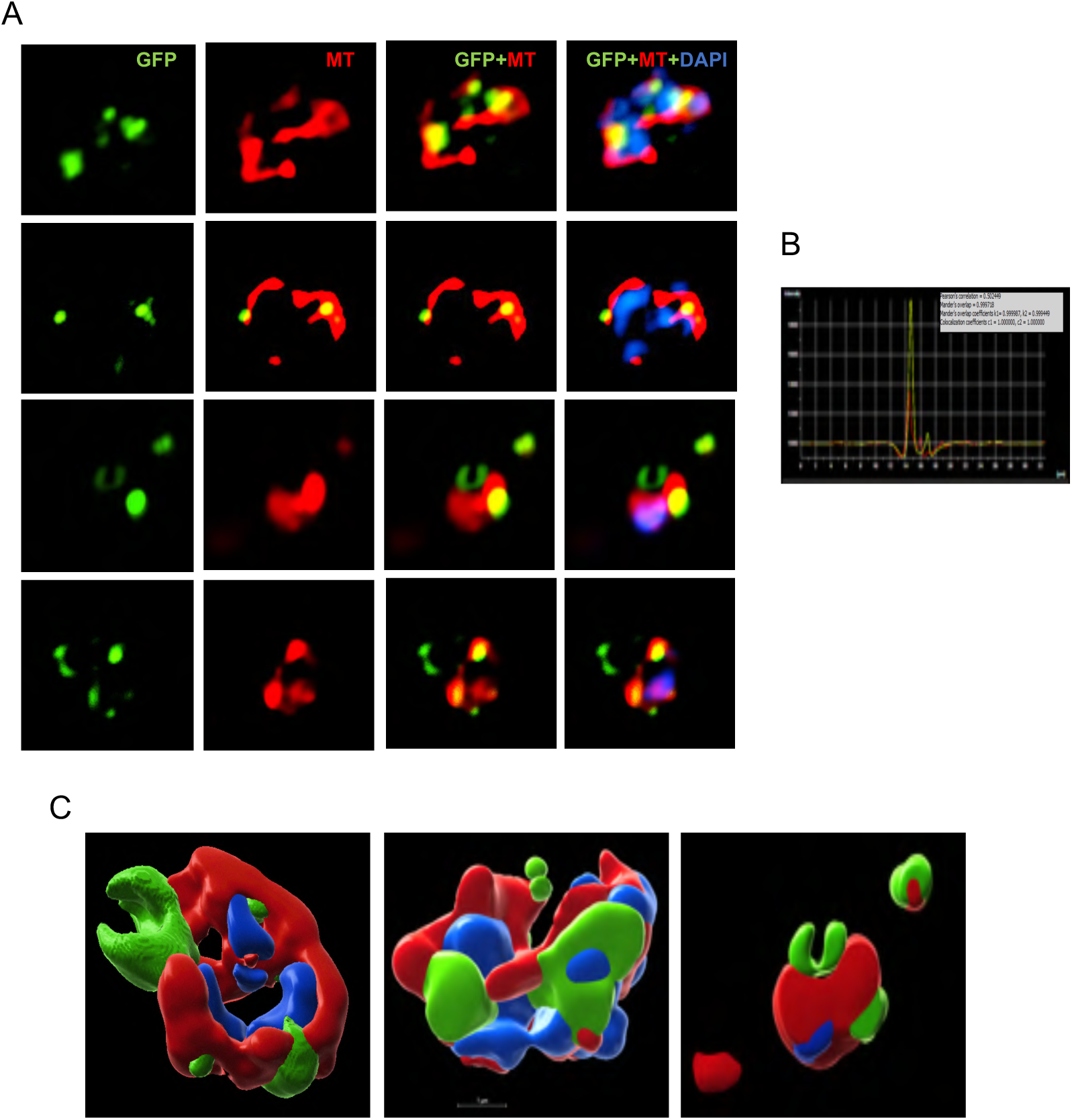
(A) SIM images of trophozoite stage transgenic parasites expressing *Pf*Dyn2-GFP fusion protein and stained with MitoTraker CMXRos dye (red), the parasite nuclei were stained with DAPI (blue). Small GFP foci of the *Pf*Dyn2-GFP protein was observed on mitochondria forming a clip-like structure at the mitochondrial putative branching point. (B) Representative histogram plot to show colocalization of mitochondria and *Pf*Dyn2-GFP tagged fusion protein in the parasite. (C) Three-dimensional reconstructions of series of Z-stack images using IMARIS software, showing co-localization of *Pf*Dyn2-GFP with mitochondrial staining of MitoTracker

Further, the transgenic parasites expressing *Pf*Dyn2-GFP were immune-stained for nuclear-encoded apicoplast proteins *Pf*ClpP. The labeling with anti-*Pf*ClpP antibody showed the apicoplast as an elongated branched structure during late-trophozoite/schizont stages, however, there was no clear overlap between GFP foci and the apicoplast (Fig. S5). Pearson’s correlation coefficient calculated for GFP labelling and apicoplast staining was found to be <0.3, which suggest that co-localization is insignificant. Similarly, the ER-labelling with ER-tracker dye did not show any clear overlap with GFP fluorescence (Fig. S6).

### Endogenous tagging of *Pf*Dyn2 with degradation domain for conditional/selective knockdown disrupt intraerythrocytic parasite developments

To study functional essentiality of *Pf*Dyn2 for parasite growth and survival, we utilized conditional/selective knockdown strategy involving FKBP destabilization domain (DD). The *pfdyn2* gene was endogenously tagged at the C-terminus with HA-DD-tag using single-crossover homologous recombination (Fig. 3), so that the fusion protein is expressed under the control of native promoter. Incorporation of C-terminal tag in the native *pfdyn2* gene was confirmed by PCR based analysis of genomic DNA from clonally selected parasites population (Fig. S7A). The integrant parasites showed expression of *Pf*Dyn2-HA-DD fusion protein of ∼95kDa, migrating at the expected predicted size for the endogenously HA-DD tagged protein. Such band could not be detected in parental 3D7 parasite line (Fig. S7B).

**Figure 3.**
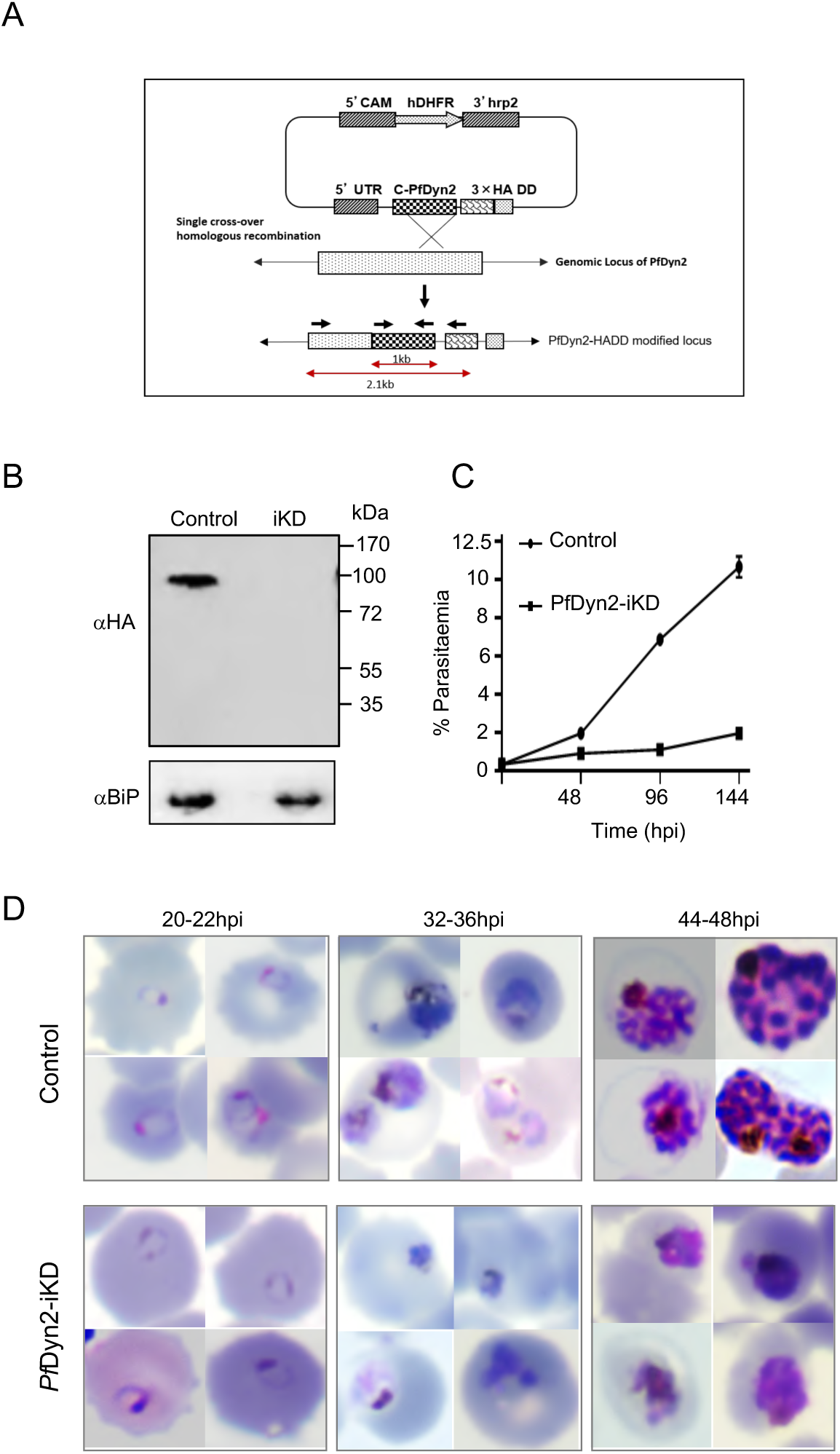
Generation of transgenic parasite expressing HA-DD tag in the *PfDyn2* gene locus, inducible knock-down of *Pf*Dyn2 protein in the transgenic parasites, and its effect on growth and development asexual cycle of the parasites. (A) Schematic representation of the strategy used to incorporate the HA-DD at the 3’ end of endogenous locus of *pfDyn2* through the single cross-over homologous recombination. (B) Western blot analysis showing reduction in the levels of fusion protein, *Pf*Dyn2-HA, in transgenic parasites grown with or without Shld1 drug (control and iKD). (C) Graph showing percentage parasitemia in *Pf*Dyn2-HA-DD transgenic parasite cultures grown with without Shld1 (control and iKD), as estimated by new ring stage parasites after each of the subsequent three developmental cycles. All analyses were performed in triplicate or more (*n*=3); the error bars indicate standard deviations. (D) Giemsa-stained images of parasites showing effect on parasite morphology at different time points (0-48h) for control and *Pf*Dyn2-iKD sets.

To study the effect of downregulation of *Pf*Dyn2 on parasite growth and development, transgenic parasites were grown in the presence or absence of Shld1 (+Shld and -Shld sets, labelled as Control and iKD respectively). To check any toxicity of Shld1 treatment, synchronous wild-type (3D7) parasites at the ring stage incubated with Shld1 ligand and calculated the parasite percentage growth for three consecutive cycles. Graph showing Shld1 is not toxic for the parasites (Fig. S7C).

The western blot analysis of parasite lysates from these two sets using anti-HA antibody showed significant reduction in levels of *Pf*Dyn2-HADD fusion protein in the *Pf*Dyn2-iKD set at 36hpi and 84hpi, thus confirming the selective degradation of HADD-tagged protein in the absence of Shld (Fig. 3B; Fig. S7D). Further, parasite growth and development were analyzed over three development cycles. In *Pf*Dyn2-iKD set, the parasite growth was found to be drastically reduced (∼90%) within three growth cycle (after 144hpi) as compared to control parasite culture (Fig. 3C). To assess the effect of *Pf*Dyn2-iKD on morphology and development, Giemsa-stained parasite smears at different time points during the asexual growth cycle were analyzed and effect on developmental stage profile were studied. As envisaged, in the control set, the parasites developed from rings to trophozoites to mature schizonts and subsequently into merozoites, which subsequently got released from the schizonts and invaded new erythrocytes, effectively increasing the total parasitemia about 5 times in each cycle (Fig. 3D). In the *Pf*Dyn2-iKD set, the parasites were able to develop from rings to trophozoites stages, however, development of the parasite from trophozoites to mature schizont was hindered, and the stressed parasites were seen as densely stained pyknotic structures (Fig. 3D). Due to these morphological and growth abnormalities, parasites were not able to develop merozoite and new ring stage parasites, causing significant reduction in overall growth of the parasites.

### Dynamin 2 knockdown disrupt mitochondrial development in the parasites

Given that *Pf*Dyn2 associates with parasite mitochondria, we assessed its functional importance in organelle segregation/fission during asexual cycle. The growth and morphology of parasite mitochondria, in control as well as in the *Pf*Dyn2-iKD set, were observed at different time points after labelling MitoTraker CMX-Ros and immune-labeling with anti-HA antibody. In the control set the mitochondria showed normal growth and development: in majority of parasites at trophozoite/late-trophozoite stages the mitochondria appeared as elongated and/or branched structure (Fig. 4A). As shown with our GFP-targeting studies, the anti-HA staining showed the *Pf*Dyn2 in distinct foci associated with branched mitochondrial at discrete points (Fig 4A.). However, the growth and development of mitochondrial was considerably affected in the *Pf*Dyn2-iKD set; majority of parasites (>80%) in *Pf*Dyn2-iKD set showed punctuate structure or diffused punctate staining of MitoTracker dye without any clear organelle structure; as expected, in the *Pf*Dyn2-iKD, there was diffused or no staining of anti-HA antibody (Fig. 4A). Based upon mitochondrial morphology and staining pattern at 33hpi, three parasite groups were identified: (i) diffusely stained/abnormal mitochondria; (ii) branched; and (iii) elongated. In the *Pf*Dyn2-iKD set majority of parasites showed diffusely stained/abnormal mitochondria (Fig. 4B and S8). In a parallel set of experiment, parasites in control and *Pf*Dyn2-iKD sets were stained with anti-HA and anti-*Pf*ClpP antibodies. In control and *Pf*Dyn2-iKD sets, the apicoplast showed elongated/branched morphology, suggesting absence of any developmental abnormalities in the apicoplast after downregulation of *Pf*Dyn2 (Fig. 4A and S9).

**Figure 4:**
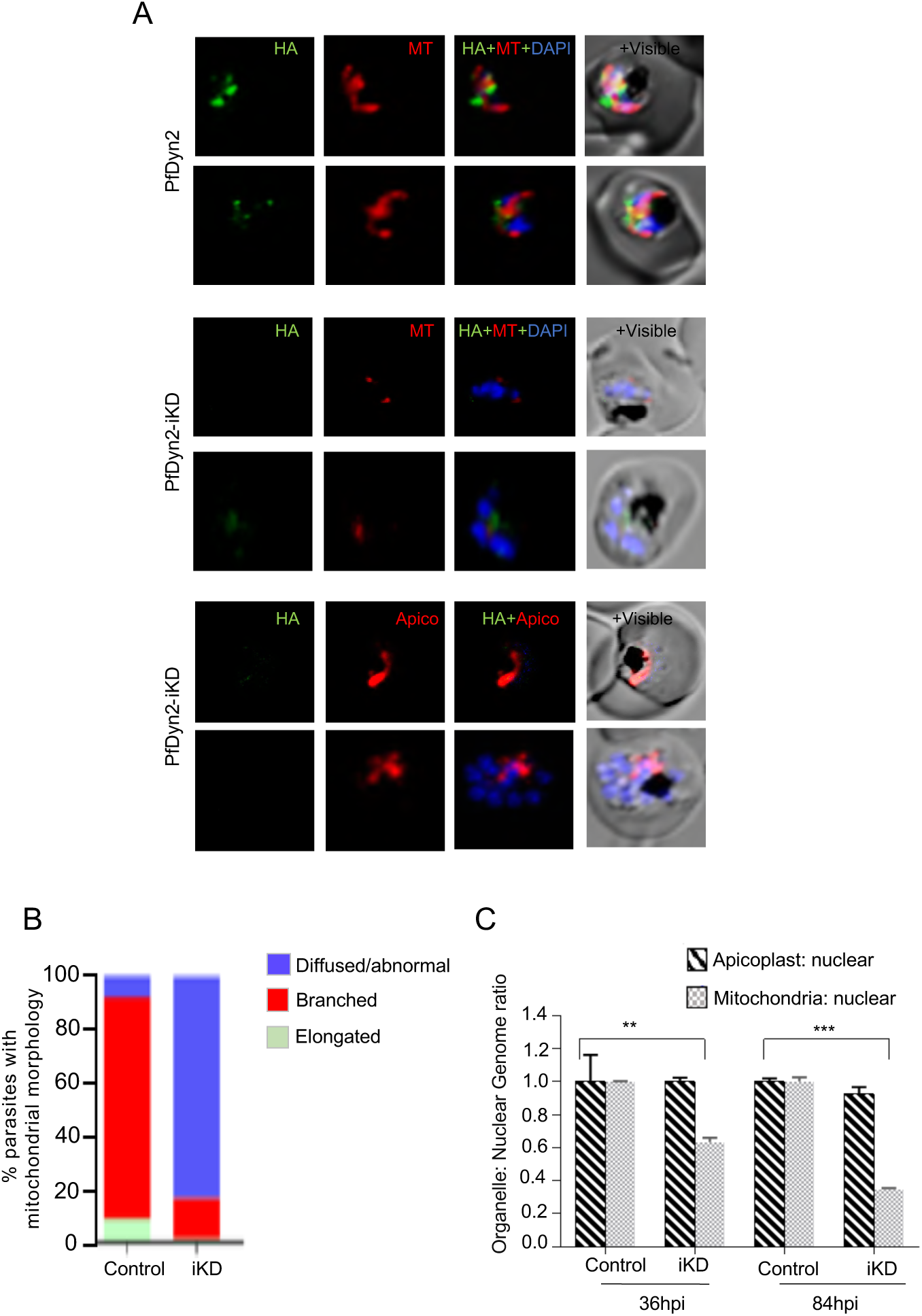
Effect of downregulation of *Pf*Dyn2 levels on growth and segregation of parasite mitochondrion. (A) Fluorescent microscopic images of transgenic *Pf*Dyn2-HA-DD parasites grown in the presence of or absence of Shld-1 drug (control and *Pf*Dyn2-iKD sets), immuno-stained with anti-HA (green) antibodies and co-stained with MitoTracker dye (red). The control parasites show elongated and branched mitochondria, whereas the iKD parasites showed deformed mitochondria at 33hpi. (B) Graph showing mitochondrial development profile in *Pf*Dyn2-iKD as compared to the control set. Based upon mitochondrial morphology and staining pattern at 33hpi, three parasite groups were identified, and percentage of parasites in each group is presented. (C) Quantitative PCR-based analysis using total DNA isolated from control and *Pf*Dyn2-iKD sets during the first and second cycle (36 and 84 hpi); graph showing normalized genomic equivalents calculated for *tufA* present on apicoplast genome, and *cox-3* present on the mitochondrial genome. Mitochondrial to nuclear genome ratio was reduced in the first and second cycle subsequently, whereas the apicoplast to nuclear genome ratio remains unaffected.

To quantitatively show this effect on mitochondrial development after selective degradation of *Pf*Dyn2 in the parasite, we assessed replication of mitochondrial genome as compared to the replication of nuclear genome in the control and *Pf*Dyn2-iKD parasite sets. The genomic equivalents of *cox-3* gene of the mitochondrial genome were determined by quantitative PCR-based analysis; in addition, the genomic equivalents of *tufA* gene of the apicoplast genome were also determined. The genomic equivalents of cox-3 in the *Pf*Dyn2-iKD set showed ∼50% reduction as compared to the control set (Fig. 4C and supplementary data), whereas minor changes were observed for *tufa* gene between control and *Pf*Dyn2-iKD sets for two consecutive cycles (Fig. 4C). These results clearly show that the down-regulation of *Pf*Dyn2 disrupts mitochondrial development and segregation but not the apicoplast in the parasites.

### Downregulation of *Pf*Dyn2 interrupts mitochondrial function and induces oxidative stress

Given that knock-down of the disrupts mitochondrial development and induce stress like phenotype with punctuate mitochondria in the parasites, we investigated effect of *Pf*Dyn2 knock-down on development of functional mitochondria and production of mitochondrial ROS in the parasite. The mitochondria membrane potential for control and *Pf*Dyn2-iKD sets was determined by using MitoProbe JC-1 staining assay. In the control set, the ratio of parasite population with mitochondrial JC-1 red and cytosolic JC-1 green staining was found to be ∼1.0, while *Pf*Dyn2-iKD showed significant reduction in the ration of JC-1 red: JC-1 green population to ∼0.5 (Fig. 5A). Our results suggesting clearly suggested loss of mitochondrial membrane potential after transient knock-down of *Pf*Dyn2

**Figure 5.**
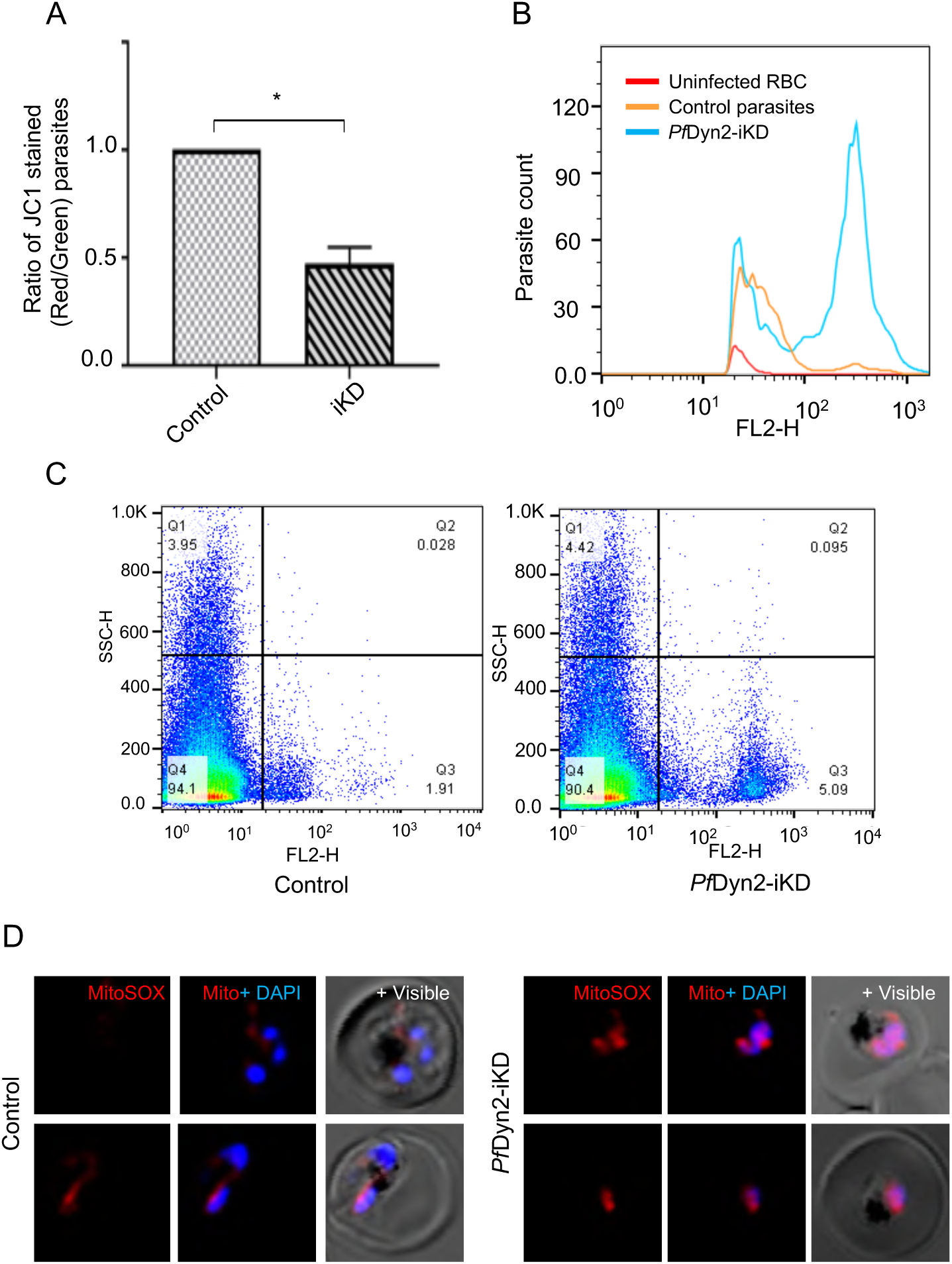
Effect of downregulation of *Pf*Dyn2 levels on mitochondrial membrane potential and mitochondrial oxidative stress. Synchronous transgenic parasites at ring trophozoite stages (33-33hpi) in control and *Pf*Dyn2-iKD sets, stained with JC1 dye or MitoSOX™ Red, Mitochondrial Superoxide Indicator, and analyzed by flow cytometry. (A) Graph showing a reduction in the ratio of JC-1 (red)/JC-1 (green) stained parasite population in the *Pf*Dyn2-iKD sets as compared to control set. (B-C) Flow cytometry histogram and dot plots showing distinct parasite population in the *Pf*Dyn2-iKD set stained with MitoSOX as compared to control set. (D) Fluorescent microscopic images of *Pf*Dyn2-iKD set stained with MitoSOX (red) dye and DAPI (blue). Parasites in the control set show diffused faint staining of elongated/branched mitochondria, whereas parasites in the *Pf*Dyn2-iKD set show intense red fluorescence in the mitochondria, suggesting enhanced levels of ROS.

The mitochondrial specific oxidative stress was assessed by using MitoSOX staining, which is an indicator for specific detection of mitochondrial superoxide in live cells. As shown in Fig. 5B-C, the control set showed a small parasite population (∼25% of parasites) with low fluorescent labeling, whereas the *Pf*Dyn2-iKD set showed a distinct parasites population with intense labelling, this set contained ∼75% of total parasites counted in the assay. Similarly, fluorescent microscopy also showed MioSOX labelling of the parasite mitochondrial in the *Pf*Dyn2-iKD set, which have punctuated mitochondria with high intensity labeling as compared to parasites in the control set having faintly stained elongated/branched mitochondria (Fig. 5D).

### Dynamin specific inhibitor block mitochondrial segregation and induce ROS production

To further confirm the role of *Pf*Dyn2 in mitochondrial fission or segregation, we assessed the effect of dynamin specific inhibitors on transgenic parasite line expressing *Pf*Dyn2-GFP. The parasites were treated at early trophozoites stages (∼27hpi) with different drugs at a concentration close to respective IC_50_ values, at mid-trophozoite stages (∼30hpi) the effect on mitochondria development and induction of mitochondrial oxidative stress was assessed. In the control set the parasite showed mitochondria with branched structure and associated *Pf*Dyn2-GFP in specific foci; whereas in the drug treated parasites, the mitochondria showed abnormal morphologies as described below. In the first set parasites were treated with dynasore, that binds the GTPase domain and inhibits GTPase activity of dynamins thus limits the mitochondrial fission (Macia et al., 2006). In dynasore treated parasites, the *Pf*Dyn2-GFP was not observed in mitochondrial associated punctate structures, only diffuse cytosolic GFP fluorescence or a single punctate GFP foci were seen (Fig. 6A). In the second set, parasites were treated with dynamin selective inhibitor Mitochondrial division inhibitor 1 (Mdivi-1), which limits the fission machinery by inhibiting Dyn2 GTPase activities. Parasites treated with Mdivi-1 showed a smaller number of GFP foci, whereas the mitochondria were present as diffusely stained or as a single punctuate structure (Fig. 6A). Based upon mitochondrial morphology and GFP localization pattern four groups of parasites were identified: (i) elongated/branched mitochondria with MitoTracker labelling and associated GFP fluorescence; (ii) fragmented/punctuate mitochondria with diffused GFP fluorescence; (iii) diffusely stained mitochondria without GFP labelling; and (iv) only GFP fluorescence without clear labelling of mitochondria. Majority of the parasite in control set showed elongated mitochondrial with associated *Pf*Dyn2-GFP punctuate fluorescence pattern as described earlier (Fig. 6B). In drug treated sets, majority of parasites showed diffusely stained or punctate mitochondria, and in some of the parasites GFP staining was not associated with the mitochondria (Fig. 6B).

**Figure 6.**
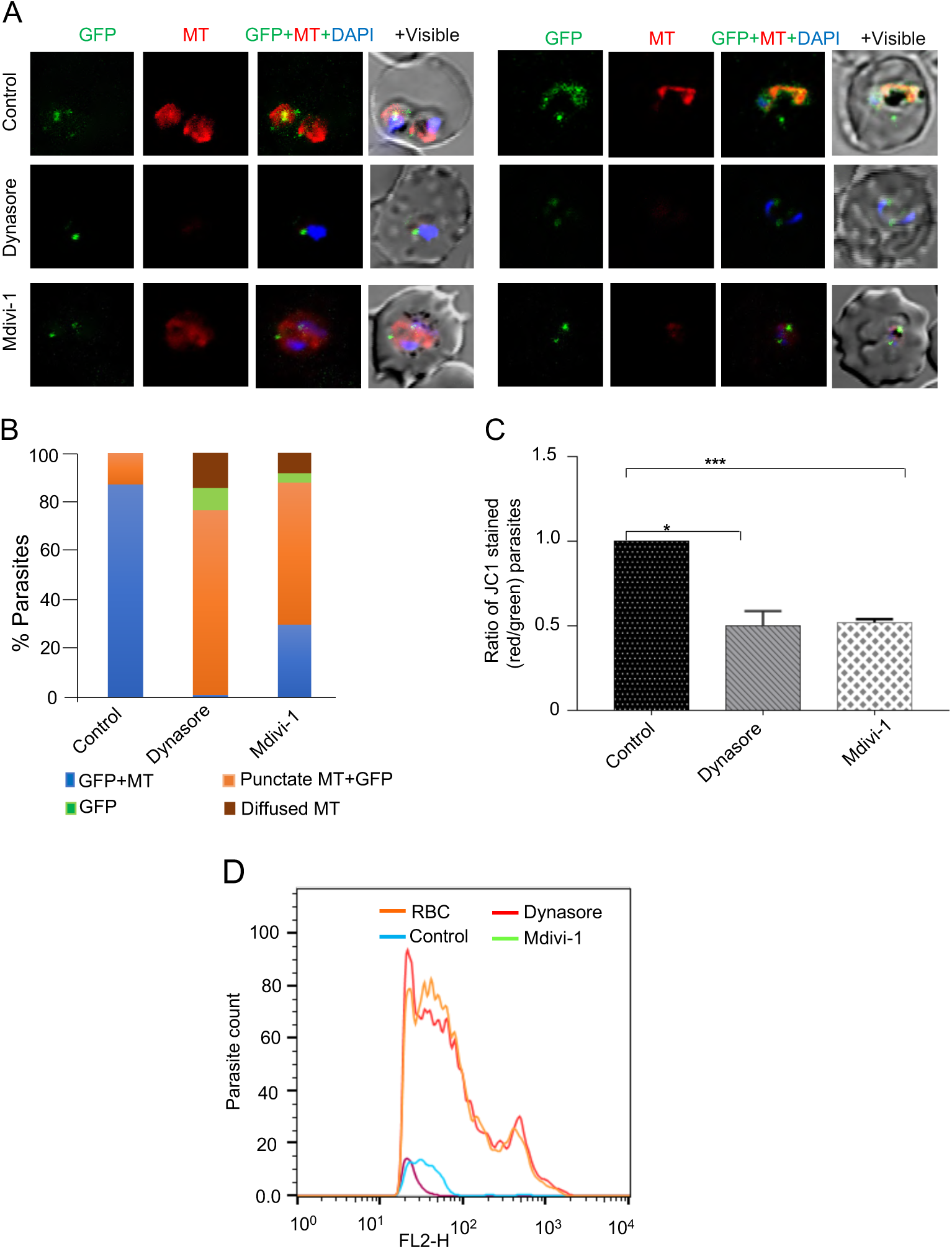
Dynamin-specific inhibitors block mitochondrial segregation, and membrane potential loss and induce ROS production: The transgenic parasites expressing *Pf*Dyn2-GFP fusion protein were treated at young trophozoite stages (27 hpi) with different drugs: dynasore (40 μM), Mdivi-1 (50 μM) or solvent (control), for 3 h. (A) Fluorescent microscopic images of parasites from drug treated and control sets stained with MitoTracker red, parasite nuclei were stained with DAPI (blue) and parasites were analysed by confocal microscopy. (B) Based upon mitochondrial morphology and GFP localization pattern, four groups were identified: (i) elongated/branched mitochondria with MitoTracker labelling and associated GFP fluorescence; (ii) fragmented/punctuate mitochondria with diffused GFP fluorescence; (iii) diffusely stained mitochondria without GFP labelling; and (iv) only GFP fluorescence without clear labelling of mitochondria. For each treatment, the percentage of parasites in each of these groups are shown. (C) The mitochondrial membrane potential (ΔΨm) levels in drug treated and control sets were measured using JC1 dye staining. Bar graph showing 40-60% reduction in the ratio of JC-1 (red)/JC-1 (green) in the treated parasites as compared to control. (D) The mitochondrial ROS levels in drug treated and control sets were measured by Mito Sox red staining. Histogram plot show a distinct parasite population in each of the drug treated sets with enhanced f Mitochondrial ROS.

The mitochondrial membrane potential was determined for each of the treatment by JC-1 assay; parasites in each of the drug treatment sets showed significant reduction in the ration of JC-1 red: JC-1 green population to ∼0.5 (Fig. 6C). These results of drug treatment on parasite mitochondrial development are similar to the data obtained with the gene knock down parasites. In parallel, for each treatment set the parasites were stained with MitoSOX red to assess mitochondrial ROS induction. As shown in Fig. 5D, treated parasites in all the three sets showed extensive staining with the MitoSOX red; the dot plot analysis of FACS data showed intense labeling in parasite populations (70-75% of parasites) all the three treatments (Fig. S10).

### C-terminal Serine S-612 phosphorylation status is not responsible for dynamin recruitment for mitochondrial fission

*Pf*Dyn2 harbor a conserved serine (Ser-612) residue in the GED region, which corresponds with the C-terminal Ser-637 in the mammalian homologue; this residue is shown to be a site for phosphorylation/dephosphorylation to influence recruitment of Dynamin to mitochondria to prevents or promotes mitochondrial fragmentation respectively [34]. To assess the role of phosphorylation/dephosphorylation status of Ser-612, we generated a transgenic parasite line overexpressing phospho-deficient mutant of *Pf*Dyn2 (S612A), labelled as *Pf*Dyn2(mut), to act as a dominant negative system; the expression levels were regulated by using ddFKBP system. Expression of HA-tagged phospho-deficient mutant *Pf*Dyn2(mut) was confirmed by immunoblotting for the parasite grown in the presence of Shld-1 drug (+Shld) whereas the control set grown in absence of Shld-1 drug (-Shld) did not show expression of the transgene (Fig. 7A). Further the effect overexpression of *Pf*Dyn2(mut) was assessed on parasites growth during three developmental cycles. However, only ∼5% parasite growth inhibition was observed in +Shld set as compare to the control set. We analyzed the subcellular localization of *Pf*Dyn2(mut) in the parasite, and effect of expression of this mutant *Pf*Dyn2 on the mitochondrial morphology and development. Immuno-labelling studies using anti-HA antibodies showed that the *Pf*Dyn2(mut) has similar localization and wild-type *Pf*Dyn2; the *Pf*Dyn2(mut) was present in discrete foci in the cytosol associated with the mitochondria (Fig. 7C). Further, there was no significant effect on the morphology and development of mitochondria in the parasites expressing *Pf*Dyn2(mut) (Fig. 7C). Similarly, there was no significant effect on the mitochondrial membrane potential due to overexpression of *Pf*Dyn2(mut) proteins (Fig. 7D). Hence our data suggesting that phosphorylation status of the Ser612 is not responsible for the membrane recruitment and functioning of the *Pf*Dyn2 in the parasite.

**Figure 7.**
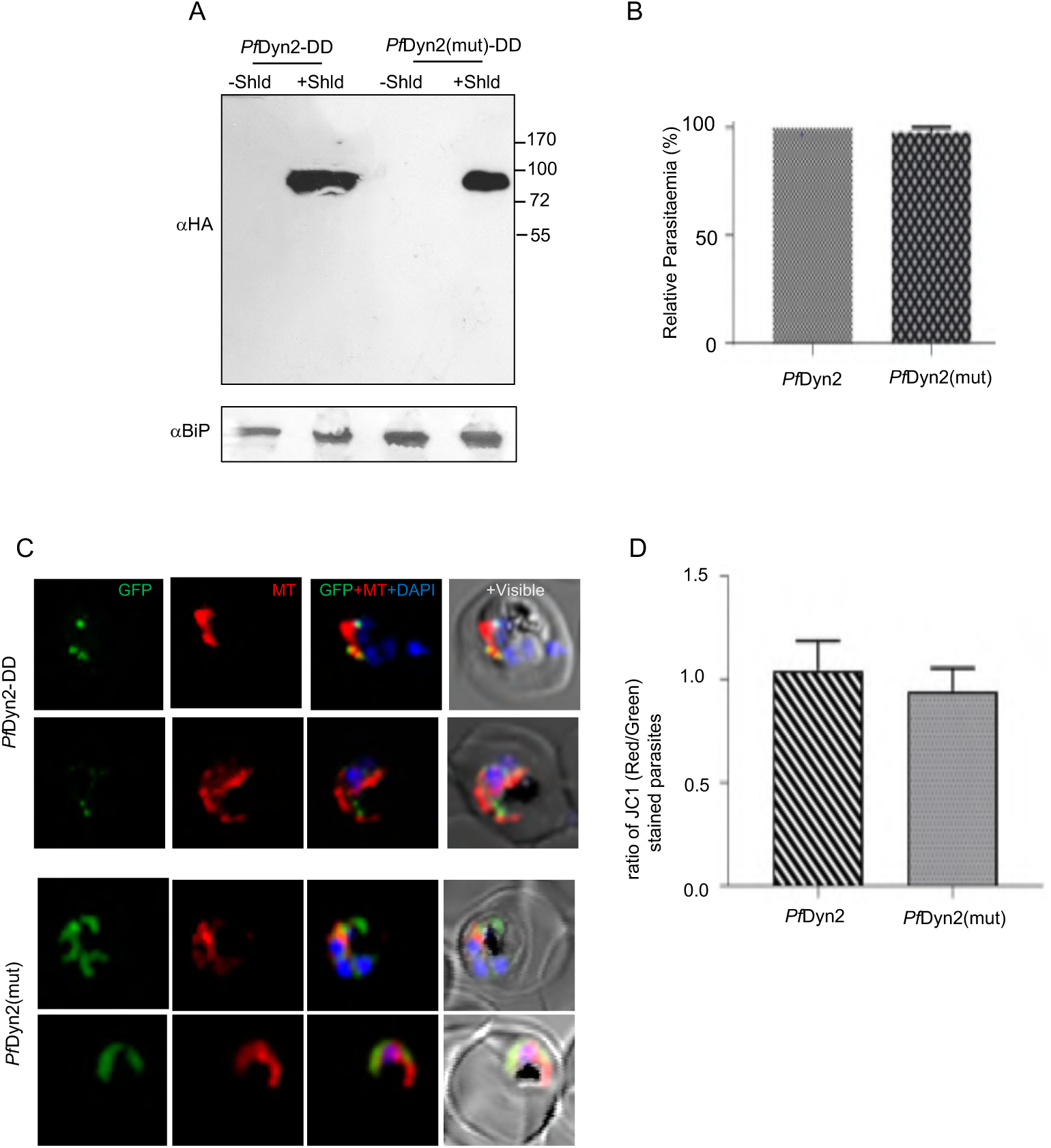
Phosphorylation/dephosphorylation of Serine(612) PfDyn2 is dispensable for dynamin recruitment and mitochondrial fission: (A) Fluorescent microscopic images of transgenic parasites expressing wild-type *Pf*Dyn2 with HA-DD tag [*Pf*Dyn2-DD], and its mutant (S612A) version [PfDyn2(mut)-DD]; the parasites were stained with MitoTracker (red) and immunolabelled with anti-HA antibody (green). Parasite nuclei were stained with DAPI and parasites were visualized by confocal microscopy. (B) Western blot analysis of lysate of *Pf*Dyn2-DD and PfDyn2(mut)-DD parasites using an anti-HA antibody. The fusion protein band (∼95 kDa) was detected in both the sets grown in presence of Shld1 drug; blot ran in parallel and probed with anti-BiP antibody was used as a loading control. (C) Graph showing parasite growth in *Pf*Dyn2(mut)-DD in the presence of Shld1 as compared to control set *Pf*Dyn2-DD growing in presence of Shld1. (D) Graph showing the ratio of JC-1 (red)/JC-1 (green) *Pf*Dyn2(mut)-DD parasites growing in the parasites with Shld1 as compared to the control set *Pf*Dyn2-DD growing in presence of Shld1.

## Discussion

Mitochondrial dynamics and metabolic pathways responsible for its genome maintenance, fission and segregation are essential for parasite survival. However, there is a poor understanding of the processes and machinery involved in organelle fission and maintenance of its synchrony along with nuclear division and segregation in the malaria parasite. Dynamin and dynamin-related protein (DRPs) are large GTPases proteins present in many eukaryotic cells, from yeast to mammals with multiple roles in protein trafficking as well as cell in organelle division [35]. One of the dynamin homologues in *P. falciparum*, *Pf*Dyn1, is shown to be involved in may be involved in the process of nutrient ingestion or endocytosis [30]. Here we have deciphered the functional role of another dynamin homologue in the parasite, *Pf*Dyn2, which phylogenetically lies in the distant cluster from *Pf*Dyn1, suggesting that is functionally different from it. The *Pf*Dyn2 harbours conserved Dynamin-N domain called as GTPase domain and C-terminus GED domain harboring conserved lysine residue, which is involved in binding of dynamins with the target membrane [36]; however, *Pf*Dyn2 lacks the PH region and PRD domain as compared to other members of the dynamin family.

In addition, *Pf*Dyn2 harbours a conserved serine (SER-612) residue in the GED region, which is shown to be a site for dephosphorylation by protein kinase A [37]. The conserved serine residue (Ser-612) in the GED region is also suggested to be a site for *Pf*Dyn2 dephosphorylation which helps in recruitment and promotes mitochondrial fission[38]. The phosphorylation and dephosphorylation of this residue have been shown to be a key regulatory mechanism for dynamin-dependent mitochondrial segregation; modification of these sites affects the mitochondrial division due to a reduction in GTPase activity [9], [38], [39], [40], [41]. Sequence analysis showed that all the critical residues, which are crucial for membrane binding and functionality are conserved in *Pf*Dyn2. Indeed, the superimposition of structural model of C-terminal domain of *Pf*Dyn2 showed strong structural similarity with structure of human DYN1 which is known to be involved in mitochondrial division (Fig. S1A).

Different members of the dynamin family are known to play diverse cellular functions, including endocytosis, organelle biogenesis, trafficking, etc. In *T. gondii*, a dynamin-related protein (TgDrpA) is shown to be associated with apicoplast [42], whereas another dynamin-related protein (TgDrpC) plays a role in mitochondrial fission [15]. Similarly, a human dynamin (Dlp1) is also known to play a role in mitochondria fission [43]. *Pf*Dyn2 showed similarities with human Dlp1, it lacks C-terminal PRD as in case of human Dlp1 [33]. To understand the potential role of *Pf*Dyn2 in the parasite, we carried out detailed localization by confocal/SIM microscopy using a transgenic parasite line expressing *Pf*Dyn2-GFP fusion protein. An earlier study, using polyclonal anti-sera for immuno-localization *Pf*Dyn2, showed partial or no clear co-localization with ER-Golgi or apicopalst in the parasite [33]. Our results with transgenic parasite lines showed that *Pf*Dyn2 [43]–[45]was localized into discrete locations associated with the mitochondria at the branching/division site at late parasite stages. In the *Plasmodium*, the mitochondrion is shown to follow a typical pattern of growth and division, which occurs in synchrony with the nuclear division and segregation[44] [29]: during the initial stages the mitochondrion is present as globular structure, it elongates during early trophozoite stages, gets branched during late-trophozoite stages just before division during schizont stages and subsequent segregation into developing daughter merozoites. The *Pf*Dyn2 is found to be associated with the branching points just before the division. Indeed, the punctate structures formed by dynamin-related proteins are known to be associated with the mitochondrial division site in other organisms [45]. The dynamins harbour self-assembling GTPase which is localized at the cytosolic surface and get associated with the outer mitochondrial membrane (OMM) as a ring/clip like structure at the sites of mitochondrial constrictions and fissions [19], [46], [47], [48], [49], this phenomenon is similar to assembly of dynamin to form a ring or spiral at the neck of a clathrin-coated pit [45], [50]. Indeed, we found that *Pf*Dyn2 is associated with the parasite mitochondrion and form a clip-like structure around the mitochondrial division site. Overall, these results suggested that *Pf*Dyn2 gets associated at mitochondrial fission site and plays role in organelle division.

To further understand functional role of *Pf*Dyn2, we utilized a transient gene knock-down strategy. A number of genetic tools have been developed for the transient down regulation of target proteins in *Plasmodium* by tagging native genes [51], which have been utilized to study the functional role of several proteins in the parasite [52], [53], [54]. Here, we have tagged the native gene at the C-terminus with a DD tag for the regulatable knock-down of the target protein in the parasite. The DD system uses a ddFKBP gene fusion and is regulated by using Shld1 drug [51], [55], [56]. Transient knock-down of *Pf*Dyn2 showed significant growth reduction and disrupted nuclear division as well as parasite development into schizonts stages. Further, this *Pf*Dyn2 knock-down also resulted in disruption of mitochondrial function and development, combined with induction of mitochondrial oxidative stress as evidenced by increase in mitochondrial ROS levels. The oxidative stress is directly linked with proper functioning of mitochondria and disruption of mitochondrial function is shown to be associated with the overproduction of ROS, which leads to mitochondrial membrane potential loss and apoptosis [57], [58] Similarly, disruption of a mitochondrial protease, regulating mitochondrial homeostasis, is also shown to induce mitochondrial oxidative stress in *P. falciparum* which leads to parasite cell death [59].

In eukaryotic cells, mitochondrial dynamics are maintained by the balanced regulation of mitochondrial fission and fusion process; suppression of mitochondrial fission or promotion of mitochondrial fusion leads to elongated forms of mitochondria. However, *Plasmodium* harbours only single mitochondria and there is no mitochondrial fusion in the parasites[60], [61] which corresponds to the absence of mitofusins in the *Plasmodium* genome. The Drp1 downregulation and upregulation have been shown to be influence mitochondrial fission and fusion process along with bidirectional relationship between mitochondrial ROS signaling. Drp1 overexpression/upregulation in certain cancer cells cause mitochondrial fragmentation, energy deficiency and increase in mitochondrial ROS [62], [63]; whereas Drp1 knock-down showed decreased mitochondrial fission [64]. In contrast, another study showed that Drp1 knockdown induces mitochondrial dysfunction, impairs mitochondrial respiration and cause upregulation of oxidative stress markers [65]. In our study as well, the data shows that disruption of mitochondrial fission by *Pf*Dyn2 knock-down leads to mitochondrial dysfunction and oxidative-stress, which ultimately caused parasite cell death.

A number of inhibitors are shown to inhibit dynamin activity by interfering with different molecular events during mitochondrial division and segregation. These inhibitors include: Dynasore, a non-competitive inhibitor of GTPase activity of dynamin; also inhibits dynamin-like GTPase involved in mitochondrial division [66]; Mdivi-1 (mitochondria division inhibitor), small-molecule inhibitor of mitochondrial fission that specifically targets Drp1[67], [68]. The association of *Pf*Dyn2 in regulating mitochondrial dynamics, function and oxidative stress were further confirmed by using these inhibitors on *Plasmodium*. Overall, our gene knock-down and inhibitor-based studies confirmed the association of *Pf*Dyn2 with mitochondrial dynamics, and its essential role in mitochondrial fission as well as in maintaining its homeostasis during asexual cycle of the parasite.

## Material and methods

### Parasite culture, plasmid constructs for transgene expression and gene knock-down, and parasite transfection

*P. falciparum* wild-type 3D7 strain and transgenic parasite lines, were cultured with 4% hematocrit in RPMI media (Invitrogen) supplemented with 0.5% Albumax using a protocol described previously. Parasite cultures were synchronized by repeated sorbitol treatment following Lambros and Vandenberg [69]. To generate a transfection vector construct, a full length *pfdyn2* (1-709 aa) was cloned in the vector pARL1a (-) vector at the *Kpn*I and *Avr*II restriction sites to express a fusion of *Pf*Dyn2 with green fluorescence protein (GFP). Synchronized *P. falciparum* 3D7 ring stage parasites were transfected with 100 μg of purified plasmid DNA (Plasmid Maxi Kit, Qiagen, Valencia, CA) by electroporation (310 V, 950 μF) [58] and the transfected parasites were selected over 2.5 nM of WR99210 drug.

To generate a transfection vector construct for C-terminal HA-DD-tagging, a C-terminal fragment of *pfdyn2* gene (1155-2130 bp) was cloned in *Xho*I and *Avr*II restriction sites of the vector pHADD [70] in frame with HA-DD cassette to yield construct pHADD-Dyn2. Transfected parasites were selected over 2.5nM WR99210 drug and 1µM Shld1; selected parasites were subsequently subjected on-and-off cycling of WR99210 drug to promote integration of the plasmid in the main genome. The transgenic parasites expressing Dyn2-HA-DD under the control of native promoter were selected by clonal selection. The parasites were constitutive grown in presence of Shld1 (Control set), whereas for transient knock-down of the target gene the parasites were grown in in absence of Shld1 (iKD-set).

To express the transgene of phosphorylation site mutant, the *pfdyn2* gene (−15 to 2130bp), with mutation in the codons of the phosphorylation site residues Ser (612) (T477G), was cloned into pHADD vector between *Xho*I and *Kpn*I restriction sites to express the transgene episomal with the HA-DD tag under the control of constitutive promoter. In another construct *pfdyn2* gene was cloned in similar manner to express the transgene wildtype protein and used as control. Transfected parasites were selected over 2.5 nM of WR99210 drug; the transgenic parasites obtained are labelled as *Pf*Dyn2(mut)-DD and *Pf*Dyn2-DD respectively. Expression of transgenes was induced by growing the cultures in presence of 1µM Shld1.

### Conditional knock-down and *in-vitro* growth assays analysis

To assess the effect of knock-down of *Pf*Dyn2 on the parasite, *Pf*Dyn2-DD transgenic parasites at tightly synchronized at ring stage were cultured in 6 well plate (0.3%–0.4% parasitemia in 2% hematocrit) and allowed to grow in media with (1µM Shld1) or without (Shld1), (+Shld and -Shld sets; control and iKD sets respectively) for three developmental cycles. Thin smears were made from each well at different time points and stained with Giemsa for microscopic analysis; parasite growth and morphology were monitored at the indicated time points. The numbers of ring/trophozoite stage parasites per 5000 RBCs were determined and percentage parasitemia [(number of infected erythrocytes/total number of erythrocytes) ×100] was calculated to assess the parasite inhibition. Each of the assays was performed three times separately on different days. To assess the effect of downregulation at protein level of *Pf*Dyn2, ring stage parasites were grown in presence of Shld1 and without Shld1 and harvested at late trophozoite stage and subjected to the immunoblotting.

For dynamin inhibitors studies, the tightly synchronized transgenic parasite culture at mid trophozoite stage (∼24-26 hpi) were treated with dynasore (80 μM), Mdivi (50 μM), or solvent (DMSO) alone for 3h.

### Isolation of parasites, differential extraction of parasite proteins and immuno-blotting

To assess the expression of *fusion protein in* transgenic *P. falciparum* parasite lines, western blot analysis was carried out. Briefly, parasites were isolated from mixed stage culture by lysis of infected erythrocyte with 0.15% saponin, the parasite pellets were suspended in Laemmli buffer, boiled, centrifuged, and the supernatant obtained was separated on 12% SDS-PAGE. The fractionated proteins were transferred from gel onto the PVDF membrane (Amersham, Piscataway, NJ, USA) and blocked-in blocking buffer (1× PBS, 0.1% Tween-20, 5% milk powder) for 2 h. The blot was washed and incubated for 1 h with primary antibody [mouse anti-GFP (1:1000); rabbit anti-BiP (1:10000); rat anti-HA (1:1000), mouse anti-spectrin (1:10000)] diluted in dilution buffer (1× PBS, 0.1% Tween-20, and 1% milk powder). Subsequently, the blot was washed and incubated for 1 h with appropriate secondary antibody (anti-rabbit, anti-rat or anti-mouse, 1:20000) conjugated to HRP, and bands were visualized by using ECL detection kit (Amersham).

### Organelle labeling, immuno-labelling, and fluorescence microscopy

*P*fDyn2-GFP transgenic parasites were synchronized by two consecutive sorbitol treatments 6 h apart. Parasites at different developmental stages were collected and stained with DAPI at a final concentration of 2 μg/ml for 10 min at 37 °C prior to imaging. To visualize the mitochondria, the *Pf*Dyn2-GFP transgenic parasites were stained with MitoTracker Red CMXRos (Invitrogen) at a final concentration of 50 nM in 1× PBS for 20 min at room temperature. Cells were washed twice with 1× PBS and subsequently fixed in 4% paraformaldehyde. The GFP expressing parasites and the parasite stained with different labeling dyes were viewed using a Nikon A1 confocal laser scanning microscope. Observations were limited to 30 min to ensure parasite viability throughout the analyses. The 3D images were constructed by using series of Z-stack images using IMARIS 7.0 (Bitplane Scientific) software. Indirect immunofluorescence assays were performed on *P. falciparum* transgenic parasite lines as described earlier [71][72]. Briefly, the parasite samples were fixed with 4% paraformaldehyde; fixed parasites were incubated for 1 h in 10% FBS in 1× PBS (blocking solution) then primary antibody [rabbit anti-*Pf*ClpP (1:200), rabbit anti-*Pf*CRT (1:500), anti-HA RAT (1:500) antibody] diluted in 10% FBS, 1× PBS added and incubated for 3h, and subsequently with Alexa-594 linked goat anti-rabbit, anti-rat or anti-mice antibody (1:500, Sigma) incubated for1 h as secondary antibody, washed thrice with 1X PBS. The parasite nuclei were stained with DAPI (2 μg/ml). The immuno-stained parasites were viewed using a Nikon A1 confocal laser scanning microscope [72].

### Mitochondria membrane potential assay

The mitochondrial membrane potential was assessed by using JC-1 staining dye as described earlier[73]. Infected erythrocytes from parasite cultures in control and experimental sets were incubated with JC-1 (5,50, 6,60 -tetrachloro-1,10, 3,30 -tetraethylbenzimidazolyl-carbocyanine iodide) at a final concentration of 5 µM, for 30 mins at 37°C. After washing with 1× PBS, the cells were examined by flow cytometry using FACS Calibur flow cytometer and CellQuestPro software (Becton Dickinson, San Jose, CA, USA), using green (488 nm) and red (635 nm) filters. Ratio of JC-1(red)/JC-1(green) was calculated to assess the loss of mitochondrial membrane potential. The JC-1-stained uninfected RBCs were used as background controls.

### Isolation of total DNA, RNA, complementary DNA synthesis, and quantitative RT-PCR

Total genomic DNA was isolated from P*f*Dyn2-HA-DD transgenic parasites from control and iKD sets (+Shld and -Shld sets respectively) at different stages following standard protocol. Gene-specific primer sets were designed using Beacon Designer 4.0 software for each organelle and nuclear genome: *tufA* (apicoplast, PF3D7_1348300), *cox3* (mitochondria, mal_mito_1), as well as for 18S rRNA (housekeeping gene as control [54]), primer details are given in Table S1. The amplification reaction contained 10ng template genomic DNA, 2× Maxima SYBR Green qPCR Master Mix (Thermo Scientific), and 10 nM gene-specific primers. Real-time PCR was performed in Micro Amp optical 96-well plates in automated ABI Step one Plus Version. Threshold cycle (Ct) values were calculated using SDS 2.4 Software (Applied Biosystem). Standard curves were used to determine genome equivalents of Ct values for respective gene and 18S ribosomal RNA for each sample. Genome equivalents of each gene were normalized using that of 18S ribosomal RNA for all the samples. For control and iKD sets (+Shld and - Shld parasite sets), the organelle: nuclear genome ratio was calculated relative to that of the control sample [72].

### Measurement of total and mitochondrial oxidative stress

To determine the Mitochondrial ROS, tightly synchronized P*f*Dyn2-HA-DD transgenic parasites culture at trophozoite stage (30-33hpi) from control and iKD sets (+Shld1 and -Shld1 respectively) were stained with MitoSOX red with 5 μM at 37 °C for 40 min in incomplete media. The stained parasites were analyzed by flow cytometry at excitation/emission wavelengths ∼510/580 nm. For microscopy, live MitoSOX-stained parasites were viewed using a Nikon A1 confocal laser scanning microscope as described above. Similarly, tightly synchronized wild-type 3D7 parasites at 30-33 hpi were treated with dynamin-specific inhibitors and stained with MitoSOX and measured by flow cytometry. Uninfected RBCs, stained in similar way, were used as background control[59].

### Statistical Analysis

The data were compared using unpaired Student’s *t*-test; the data sets were analyzed, and the graphical presentations were made using GraphPad Prism ver 5.0.

## Data Availability

All relevant data are within the manuscript and its Supporting Information files.

## Competing interests

The authors declare that they have no competing interests

## Acknowledgements

We are grateful to Tim Gilberger for providing the vector pARL1a(-), Alan cowman and Alex Maier for providing the pHADD vector. We thank Rotary blood bank, New Delhi for providing RBCs.

## Funding

The research work in AM’s laboratory is supported by Flagship Grant (#RAD-40/104/2023-MED-DBT) and Centre of Excellence grant (BT/COE/ 34/SP15138/2015) from the Department of Biotechnology (Govt. of India), and Research grant (#CRG/2022/002655) from the Science and Engineering Research Board, Department of Science and Technology (Govt. of India). VT is supported by BioCARe research grant (#BT/PR19127/BIC/101/1047/2016) from the Department of Biotechnology, Govt. of India; AM and PA are supported by a research fellowship from the Department of Biotechnology, Govt. of India; MEH was supported by Pre-doctoral fellowship from ICGEB. The funders had no role in study design, data collection and analysis, decision to publish, or preparation of the manuscript.

## Authors’ Contributions

AM conceived and designed the study; VT, SS, SR, AzM, MMB, PA, MEH and SJ carried out the experiments; AM, and SA analyzed the data; and AM and VT wrote the paper with contributions from all the authors.

## Notes

### Competing Interest Statement

The authors have declared no competing interest.

## References

[1] World Health Organization, 20 years of global progress and challenges, vol. WHO/HTM/GM, no. December. 2020.

[2] J. Hemingway et al., “Tools and Strategies for Malaria Control and Elimination: What Do We Need to Achieve a Grand Convergence in Malaria?,” PLoS Biol, vol. 14, no. 3, 2016, doi: 10.1371/journal.pbio.1002380.

[3] A. Owusu-Ofori, D. Gadzo, and I. Bates, “Transfusion-transmitted malaria: donor prevalence of parasitaemia and a survey of healthcare workers knowledge and practices in a district hospital in Ghana,” Malar J, vol. 15, p. 234, 2016, doi: 10.1186/s12936-016-1289-3.

[4] L. Tilokani, S. Nagashima, V. Paupe, and J. Prudent, “Mitochondrial dynamics: Overview of molecular mechanisms,” Essays in Biochemistry, vol. 62, no. 3. pp. 341–360, 2018. doi: 10.1042/EBC20170104.

[5] K. J. Kamer and V. K. Mootha, “The molecular era of the mitochondrial calcium uniporter,” Nature Reviews Molecular Cell Biology, vol. 16, no. 9. 2015. doi: 10.1038/nrm4039.

[6] L. Tilokani, S. Nagashima, V. Paupe, and J. Prudent, “Mitochondrial dynamics: Overview of molecular mechanisms,” Essays in Biochemistry, vol. 62, no. 3. 2018. doi: 10.1042/EBC20170104.

[7] I. Scott and R. J. Youle, “Mitochondrial fission and fusion,” Essays Biochem, vol. 47, p. 85, 2010, doi: 10.1042/BSE0470085.

[8] H. Chen and D. C. Chan, “Mitochondrial dynamics-fusion, fission, movement, and mitophagy-in neurodegenerative diseases,” Hum Mol Genet, vol. 18, no. R2, 2009, doi: 10.1093/hmg/ddp326.

[9] C. R. Chang and C. Blackstone, “Dynamic regulation of mitochondrial fission through modification of the dynamin-related protein Drp1,” in *Annals of the New York Academy of Sciences*, Ann N Y Acad Sci, 2010, pp. 34–39. doi: 10.1111/j.1749-6632.2010.05629.x.

[10] S. Frank et al., “The Role of Dynamin-Related Protein 1, a Mediator of Mitochondrial Fission, in Apoptosis,” Dev Cell, 2001, doi: 10.1016/S1534-5807(01)00055-7.

[11] Y. Fannjiang et al., “Mitochondrial fission proteins regulate programmed cell death in yeast.,” Genes Dev, 2004, doi: 10.1101/gad.1247904.

[12] R. Kalia et al., “Structural basis of mitochondrial receptor binding and constriction by DRP1,” Nature 2018 558:7710, vol. 558, no. 7710, pp. 401–405, Jun. 2018, doi: 10.1038/s41586-018-0211-2.

[13] A. D. Mozdy, J. M. Mccaffery, and J. M. Shaw, “Dnm1p GTPase-mediated Mitochondrial Fission Is a Multi-step Process Requiring the Novel Integral Membrane Component Fis1p,” 2000. [Online]. Available: http://www.jcb.org/cgi/content/full/151/2/367

[14] S. C. J. Helle et al., “Mechanical force induces mitochondrial fission,” Elife, vol. 6, 2017, doi: 10.7554/eLife.30292.

[15] C. Melatti et al., “A unique dynamin-related protein is essential for mitochondrial fission in Toxoplasma gondii,” 2019, doi: 10.1371/journal.ppat.1007512.

[16] C. A. Francy, F. J. D. Alvarez, L. Zhou, R. Ramachandran, and J. A. Mears, “The mechanoenzymatic core of dynamin-related protein 1 comprises the minimal machinery required for membrane constriction,” Journal of Biological Chemistry, vol. 290, no. 18, pp. 11692–11703, May 2015.

[17] D. Otsuga et al., “The dynamin-related GTPase, Dnm1p, controls mitochondrial morphology in yeast,” Journal of Cell Biology, vol. 143, no. 2, pp. 333–349, 1998, doi: 10.1083/jcb.143.2.333.

[18] E. Smirnova, D. L. Shurland, S. N. Ryazantsev, and A. M. Van Der Bliek, “A human dynamin-related protein controls the distribution of mitochondria,” Journal of Cell Biology, vol. 143, no. 2, pp. 351–358, 1998, doi: 10.1083/jcb.143.2.351.

[19] S. I. Arimura and N. Tsutsumi, “A dynamin-like protein (ADL2b), rather than FtsZ, is involved in Arabidopsis mitochondrial division,” Proc Natl Acad Sci U S A, vol. 99, no. 8, pp. 5727–5731, 2002, doi: 10.1073/pnas.082663299.

[20] E. B. Gkerküçük, M. Tramier, and G. Bertolin, “Imaging mitochondrial functions: From fluorescent dyes to genetically-encoded sensors,” Genes, vol. 11, no. 2. 2020. doi: 10.3390/genes11020125.

[21] C. Y. Guo, L. Sun, X. P. Chen, and D. S. Zhang, “Oxidative stress, mitochondrial damage and neurodegenerative diseases,” Neural Regen Res, vol. 8, no. 21, pp. 2003–2014, 2013, doi: 10.3969/j.issn.1673-5374.2013.21.009.

[22] D. B. Zorov, M. Juhaszova, and S. J. Sollott, “Mitochondrial reactive oxygen species (ROS) and ROS-induced ROS release,” Physiological Reviews, vol. 94, no. 3. pp. 909– 950, 2014. doi: 10.1152/physrev.00026.2013.

[23] R. Yu et al., “The phosphorylation status of Ser-637 in dynamin-related protein 1 (Drp1) does not determine Drp1 recruitment to mitochondria,” Journal of Biological Chemistry, vol. 294, no. 46, pp. 17262–17277, 2019, doi: 10.1074/jbc.RA119.008202.

[24] B. Cho et al., “CDK5-dependent inhibitory phosphorylation of Drp1 during neuronal maturation,” Exp Mol Med, vol. 46, no. 7, p. e105, 2014, doi: 10.1038/EMM.2014.36.

[25] “Evidence for Golgi-independent transport from the early secretory pathway to the plastid in malaria parasites - Tonkin - 2006 - Molecular Microbiology - Wiley Online Library.” Accessed: Mar. 30, 2023. [Online]. Available: https://onlinelibrary.wiley.com/doi/10.1111/j.1365-2958.2006.05244.x

[26] C. J. Tonkin, N. S. Struck, K. A. Mullin, L. M. Stimmler, and G. I. McFadden, “Evidence for Golgi-independent transport from the early secretory pathway to the plastid in malaria parasites,” Mol Microbiol, vol. 61, no. 3, pp. 614–630, Aug. 2006, doi: 10.1111/J.1365-2958.2006.05244.X.

[27] C. Tomova, B. M. Humbel, W. J. C. Geerts, R. Entzeroth, J. C. M. Holthuis, and A. J. Verkleij, “Membrane contact sites between apicoplast and ER in Toxoplasma gondii revealed by electron tomography,” Traffic, vol. 10, no. 10, 2009, doi: 10.1111/j.1600-0854.2009.00954.x.

[28] M. Nishi, K. Hu, J. M. Murray, and D. S. Roos, “Organellar dynamics during the cell cycle of Toxoplasma gondii,” J Cell Sci, vol. 121, no. Pt 9, p. 1559, May 2008, doi: 10.1242/JCS.021089.

[29] M. O. Anwar, M. M. Islam, V. Thakur, I. kaur, and A. Mohmmed, “Defining ER-mitochondria contact dynamics in Plasmodium falciparum by targeting component of phospholipid synthesis pathway, phosphatidylserine synthase (PfPSS),” Mitochondrion, vol. 65, pp. 124–138, Jul. 2022, doi: 10.1016/J.MITO.2022.05.005.

[30] H. Li et al., “Isolation and functional characterization of a dynamin-like gene from Plasmodium falciparum,” Biochem Biophys Res Commun, vol. 320, no. 3, pp. 664–671, Jul. 2004, doi: 10.1016/J.BBRC.2004.06.010.

[31] H. C. Zhou, Y. H. Gao, X. Zhong, and H. Wang, “Dynamin like protein 1 participated in the hemoglobin uptake pathway of Plasmodium falciparum,” Chin Med J (Engl), vol. 122, no. 14, pp. 1686–1691, Jul. 2009, doi: 10.3760/CMA.J.ISSN.0366-6999.2009.14.015.

[32] R. Jilly, N. Z. Khan, H. Aronsson, and D. Schneider, “Dynamin-like proteins are potentially involved in membrane dynamics within chloroplasts and cyanobacteria,” Front Plant Sci, vol. 9, p. 22, Feb. 2018, doi: 10.3389/FPLS.2018.00206/FULL.

[33] S. Charneau et al., “Characterization of PfDYN2, a dynamin-like protein of Plasmodium falciparum expressed in schizonts,” Microbes Infect, vol. 9, no. 7, pp. 797–805, Jun. 2007, doi: 10.1016/J.MICINF.2007.02.020.

[34] R. Kandimalla and P. H. Reddy, “Multiple faces of dynamin-related protein 1 and its role in Alzheimer’s disease pathogenesis,” Biochimica et Biophysica Acta - Molecular Basis of Disease. 2016. doi: 10.1016/j.bbadis.2015.12.018.

[35] C. A. Konopka, J. B. Schleede, A. R. Skop, and S. Y. Bednarek, “Dynamin and Cytokinesis,” Traffic, vol. 7, no. 3, pp. 239–247, Mar. 2006, doi: 10.1111/J.1600-0854.2006.00385.X.

[36] H. H. Niemann, M. L. W. Knetsch, A. Scherer, D. J. Manstein, and F. J. Kull, “Crystal structure of a dynamin GTPase domain in both nucleotide-free and GDP-bound forms,” EMBO J, vol. 20, no. 21, pp. 5813–5821, Nov. 2001, doi: 10.1093/EMBOJ/20.21.5813.

[37] C. R. Chang and C. Blackstone, “Dynamic regulation of mitochondrial fission through modification of the dynamin-related protein Drp1,” Ann N Y Acad Sci, vol. 1201, no. 1, pp. 34–39, Jul. 2010, doi: 10.1111/J.1749-6632.2010.05629.X.

[38] J. T. Cribbs and S. Strack, “Reversible phosphorylation of Drp1 by cyclic AMP-dependent protein kinase and calcineurin regulates mitochondrial fission and cell death,” EMBO Rep, vol. 8, no. 10, p. 939, Oct. 2007, doi: 10.1038/SJ.EMBOR.7401062.

[39] C. R. Chang and C. Blackstone, “Cyclic AMP-dependent protein kinase phosphorylation of Drp1 regulates its GTPase activity and mitochondrial morphology,” J Biol Chem, vol. 282, no. 30, pp. 21583–21587, Jul. 2007, doi: 10.1074/JBC.C700083200.

[40] C. H. Chou et al., “GSK3beta-Mediated Drp1 Phosphorylation Induced Elongated Mitochondrial Morphology against Oxidative Stress,” PLoS One, vol. 7, no. 11, 2012, doi: 10.1371/JOURNAL.PONE.0049112.

[41] R. Kandimalla and P. H. Reddy, “Multiple faces of dynamin-related protein 1 and its role in Alzheimer’s disease pathogenesis,” Biochim Biophys Acta, vol. 1862, no. 4, pp. 814–828, Apr. 2016, doi: 10.1016/J.BBADIS.2015.12.018.

[42] G. G. van Dooren, S. B. Reiff, C. Tomova, M. Meissner, B. M. Humbel, and B. Striepen, “A Novel Dynamin-Related Protein Has Been Recruited for Apicoplast Fission in Toxoplasma gondii,” Current Biology, vol. 19, no. 4, 2009, doi: 10.1016/j.cub.2008.12.048.

[43] E. Smirnova, L. Griparic, D.-L. Shurland, and A. M. van der Bliek, “Dynamin-related Protein Drp1 Is Required for Mitochondrial Division in Mammalian Cells,” Mol Biol Cell, 2001, doi: 10.1091/mbc.12.8.2245.

[44] G. G. Van Dooren, M. Marti, C. J. Tonkin, L. M. Stimmler, A. F. Cowman, and G. I. McFadden, “Development of the endoplasmic reticulum, mitochondrion and apicoplast during the asexual life cycle of Plasmodium falciparum,” Mol Microbiol, vol. 57, no. 2, pp. 405–419, Jul. 2005, doi: 10.1111/J.1365-2958.2005.04699.X.

[45] S. Y. Miyagishima et al., “A Plant-Specific Dynamin-Related Protein Forms a Ring at the Chloroplast Division Site,” Plant Cell, vol. 15, no. 3, pp. 655–665, Mar. 2003, doi: 10.1105/TPC.009373.

[46] A. M. Labrousse, M. D. Zappaterra, D. A. Rube, and A. M. Van der Bliek, “C. elegans Dynamin-Related Protein DRP-1 Controls Severing of the Mitochondrial Outer Membrane,” Mol Cell, vol. 4, no. 5, pp. 815–826, Nov. 1999, doi: 10.1016/S1097-2765(00)80391-3.

[47] W. Bleazard et al., “The dynamin-related GTPase Dnm1 regulates mitochondrial fission in yeast,” Nat Cell Biol, vol. 1, no. 5, pp. 298–304, 1999, doi: 10.1038/13014.

[48] A. M. Labrousse, M. D. Zappaterra, D. A. Rube, and A. M. van der Bliek, “C. elegans dynamin-related protein DRP-1 controls severing of the mitochondrial outer membrane.,” Mol Cell, vol. 4, no. 5, pp. 815–826, 1999, doi: 10.1016/s1097-2765(00)80391-3.

[49] W. Bleazard et al., “The dynamin-related GTPase Dnm1 regulates mitochondrial fission in yeast,” Nat Cell Biol, 1999, doi: 10.1038/13014.

[50] J. E. Hinshaw and S. L. Schmid, “Dynamin self-assembles into rings suggesting a mechanism for coated vesicle budding,” Nature, 1995, doi: 10.1038/374190a0.

[51] C. M. Armstrong and D. E. Goldberg, “An FKBP destabilization domain modulates protein levels in Plasmodium falciparum,” Nat Methods, vol. 4, no. 12, pp. 1007–1009, Dec. 2007, doi: 10.1038/NMETH1132.

[52] S. Jain et al., “The prokaryotic ClpQ protease plays a key role in growth and development of mitochondria in Plasmodium falciparum,” Cell Microbiol, vol. 15, no. 10, pp. 1660–1673, Oct. 2013, doi: 10.1111/CMI.12142/SUPPINFO.

[53] P. K. Sheokand, M. Narwal, V. Thakur, and A. Mohmmed, “GlmS mediated knock-down of a phospholipase expedite alternate pathway to generate phosphocholine required for phosphatidylcholine synthesis in Plasmodium falciparum,” Biochemical Journal, vol. 478, no. 18, pp. 3429–3444, Sep. 2021, doi: 10.1042/BCJ20200549.

[54] G. Datta, M. E. Hossain, M. Asad, S. Rathore, and A. Mohmmed, “Plasmodium falciparum OTU-like cysteine protease (PfOTU) is essential for apicoplast homeostasis and associates with noncanonical role of Atg8,” Cell Microbiol, vol. 19, no. 9, p. e12748, Sep. 2017, doi: 10.1111/CMI.12748.

[55] J. D. Dvorin, A. K. Bei, B. I. Coleman, and M. T. Duraisingh, “Functional diversification between two related Plasmodium falciparum merozoite invasion ligands is determined by changes in the cytoplasmic domain,” Mol Microbiol, vol. 75, no. 4, p. 990, Feb. 2010, doi: 10.1111/J.1365-2958.2009.07040.X.

[56] G. J. Hermann et al., “Mitochondrial fusion in yeast requires the transmembrane GTPase Fzo1p,” Journal of Cell Biology, 1998, doi: 10.1083/jcb.143.2.359.

[57] D. B. Zorov, M. Juhaszova, and S. J. Sollott, “Mitochondrial reactive oxygen species (ROS) and ROS-induced ROS release,” Physiol Rev, vol. 94, no. 3, pp. 909–950, Jul. 2014, doi: 10.1152/PHYSREV.00026.2013.

[58] S. Marchi et al., “Mitochondria-ros crosstalk in the control of cell death and aging,” J Signal Transduct, vol. 2012, pp. 1–17, Nov. 2012, doi: 10.1155/2012/329635.

[59] S. Singh et al., “Dual role of an essential HtrA2/Omi protease in the human malaria parasite: Maintenance of mitochondrial homeostasis and induction of apoptosis-like cell death under cellular stress,” PLoS Pathog, vol. 18, no. 10, p. e1010932, Oct. 2022, doi: 10.1371/JOURNAL.PPAT.1010932.

[60] R. R. Stanway et al., “Organelle segregation into Plasmodium liver stage merozoites,” 2011, doi: 10.1111/j.1462-5822.2011.01657.x.

[61] L. Voleman and P. Doležal, “Mitochondrial dynamics in parasitic protists Mitochondrial dynamics in model cellular system of yeast and humans,” 2019, doi: 10.1371/journal.ppat.1008008.

[62] Q. Huang et al., “Increased mitochondrial fission promotes autophagy and hepatocellular carcinoma cell survival through the ROS-modulated coordinated regulation of the NFKB and TP53 pathways,” Autophagy, vol. 12, no. 6, pp. 999–1014, Jun. 2016, doi: 10.1080/15548627.2016.1166318/SUPPL_FILE/KAUP_A_1166318_SM3342.DOCX.

[63] Z. Huang et al., “Sorting Nexin 5 Plays an Important Role in Promoting Ferroptosis in Parkinson’s Disease,” Oxid Med Cell Longev, vol. 2022, 2022, doi: 10.1155/2022/5463134.

[64] D. Kim, A. Sankaramoorthy, and S. Roy, “Downregulation of Drp1 and Fis1 Inhibits Mitochondrial Fission and Prevents High Glucose-Induced Apoptosis in Retinal Endothelial Cells,” Cells, vol. 9, no. 7, Jul. 2020, doi: 10.3390/CELLS9071662.

[65] M. Dulac et al., “Drp1 knockdown induces severe muscle atrophy and remodelling, mitochondrial dysfunction, autophagy impairment and denervation,” J Physiol, vol. 598, no. 17, pp. 3691–3710, Sep. 2020, doi: 10.1113/JP279802.

[66] E. Macia, M. Ehrlich, and R. H. Massol, “Dynasore, a Cell-Permeable Inhibitor of Dynamin”, doi: 10.1016/j.devcel.2006.04.002.

[67] Y. Deng, S. Li, Z. Chen, W. Wang, B. Geng, and J. Cai, “Mdivi-1, a mitochondrial fission inhibitor, reduces angiotensin-II-induced hypertension by mediating VSMC phenotypic switch,” Biomedicine & Pharmacotherapy, vol. 140, p. 111689, Aug. 2021, doi: 10.1016/J.BIOPHA.2021.111689.

[68] Y. H. Li et al., “Mdivi-1, a mitochondrial fission inhibitor, modulates T helper cells and suppresses the development of experimental autoimmune encephalomyelitis,” J Neuroinflammation, vol. 16, no. 1, pp. 1–11, Jul. 2019, doi: 10.1186/S12974-019-1542-0/FIGURES/7.

[69] C. Lambros and J. P. Vanderberg, “Synchronization of *Plasmodium falciparum* erythrocytic stages in culture,” J Parasitol, vol. 65, no. 3, pp. 418–420, Jun. 1979.

[70] B. S. Crabb et al., “Transfection of the human malaria parasite Plasmodium falciparum.,” Methods Mol Biol, vol. 270, pp. 263–276, 2004, doi: 10.1385/1-59259-793-9:263.

[71] T. Wickramarachchi, Y. S. Devi, A. Mohmmed, and V. S. Chauhan, “Identification and characterization of a novel *Plasmodium falciparum* merozoite apical protein involved in erythrocyte binding and invasion,” PLoS One, vol. 3, no. 3, p. e1732, Mar. 2008, doi: 10.1371/journal.pone.0001732.

[72] G. Datta, M. E. Hossain, M. Asad, S. Rathore, and A. Mohmmed, “Plasmodium falciparum OTU-like cysteine protease (PfOTU) is essential for apicoplast homeostasis and associates with noncanonical role of Atg8,” Cell Microbiol, vol. 19, no. 9, 2017, doi: 10.1111/cmi.12748.

[73] S. Rathore et al., “Disruption of a mitochondrial protease machinery in Plasmodium falciparum is an intrinsic signal for parasite cell death,” Cell Death Dis, vol. 2, no. 11, Nov. 2011, doi: 10.1038/CDDIS.2011.118.

