## Supplementary for "A dynamin-like protein in the human malaria parasite plays an essential role in mitochondrial homeostasis and fission during asexual blood stages"

**Supplementary Data:**

**Sequence analysis and targeting prediction of *Pf*DYN**

The *P. falciparum* Dyn2 (*Pf*Dyn2) is a 709 amino acid long protein; multiple sequence alignment of PfDYN2 with DYN proteins from other organisms showed that it is highly conserved in *Arabidopsis thaliana* (41% identical, E-value 1e-136) and lowest in *Homo sapiens* (34% identical E-value 5e-136). The C-terminal region is highly conserved, which comprises conserved lysine residue responsible for membrane binding. There is a conserved serine residue that has been shown as a dynamin phosphorylation or de-phosphorylation site with the regulatory subunit of phosphatidylinositol 3-kinase (PKC) is shown (Figure S1A, B). To illustrate the structural homology of *Pf*DYN2 with the template structure, overlapping structure models are shown for the C-terminal region was done by Phyre2 and analyzed by Chimera 1.2 software. (Figure S1A). The C-terminal *Pf*DYN2 covers 101 residues (51% of C-terminal sequence with RMSD value 0. 003Å) as compared with the human dynamin 1 as template model. The RMSD value less than 0.2Å indicated the best similarly or conservation. In addition, the C-terminal region was found to be highly conserved among different species of *Plasmodium* as well as other organisms (Fig. S1 B).

**Drug Revival Assay**

To check drug cytotoxicity, the tightly synchronized transgenic parasite culture at mid trophozoite stage (~24-26 hpi) was treated with dynasore (80 μM), Mdivi (50 μM) or solvent (DMSO) alone for 3h and washed off with twice 1X PBS. Further, fresh media is supplemented and incubated at 37º C, and calculated formation of rings.

**Table S1: List of primers used in the study:**

| Primer | Sequence |
|---|---|
| 1078A | GAATTCGGATCCGTTTTACCAGTGATTCCAG |
| 1079A | CTCGAGGTCGACAACTTCTTGGTTTCTAATTTC |
| 1606A | CAAAATGCTTAAGACAGATCTTCGG |
| D1 | GCCCGCGGTTCACCATTAGAAATGTTAACAGATG |
| D2 | GCGGTACCAACTTCTTGGTTTCTAATTTCGGC |
| Cox3 (FP) | ATATGATACTTCTACCGAA |
| cox3 (RP) | CCAGATTATTTCAACAAAA |
| 18S rRNA (FP) | GCTGACTACGTCCCTGCCC |
| 18S rRNA (RP) | ACAATTCATCATATCTTTCAATCGGTA |
| tufA (FP) | GATATTGATTCAGCTCCAGAAGAAA |
| tufA (RP) | ATATCCATTTGTGTGGCTCCTATAA |

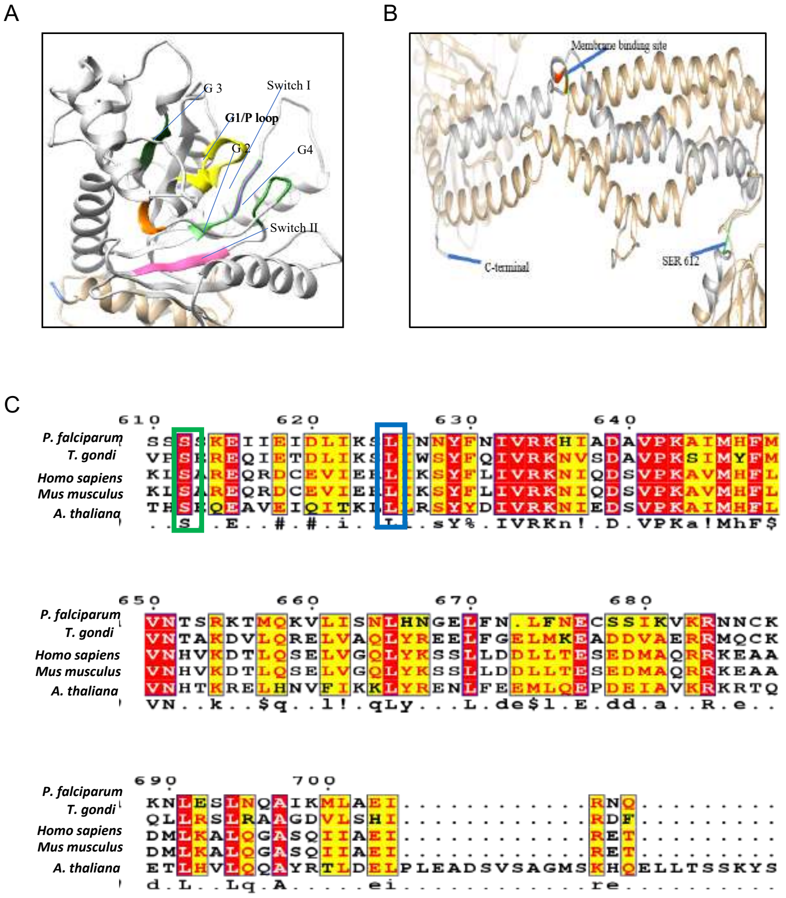


Figure S1

**Figure S1.** Predicted 3D structure of N-terminal *Pf*Dyn2 and sequence homology. (A) The The structure model of *Pf*Dyn2: white colour ribbon shows Dynamin-N-domain/Dlp1 region; Yellow colour is representing conserved motif GXXXGS/T; the four G1-G4 region conserved are labelled and two switches are marked in pink. (B) The structure model of C-terminal GED domain of *Pf*Dyn2 showing the membrane binding region (red). Conserved Serine (Serine-612) is marked in green, which is phosphorylation/dephosphorylation site crucial for the functional role of dynamin in other model organisms. (C) Amino acid sequence alignment of GED region of *Pf*Dyn2 with that of Dynamin homologs from *Mus musculus*, *Homo sapiens*, *Toxoplasma gondi*, and *Arabidopsis thaliana*. Amino acids with ≥ 60% identity are shown in pink box, the conserved lysine residue critical for membrane recruitment of dynamin is marked with blue box, and conserved serine residue is marked in green.

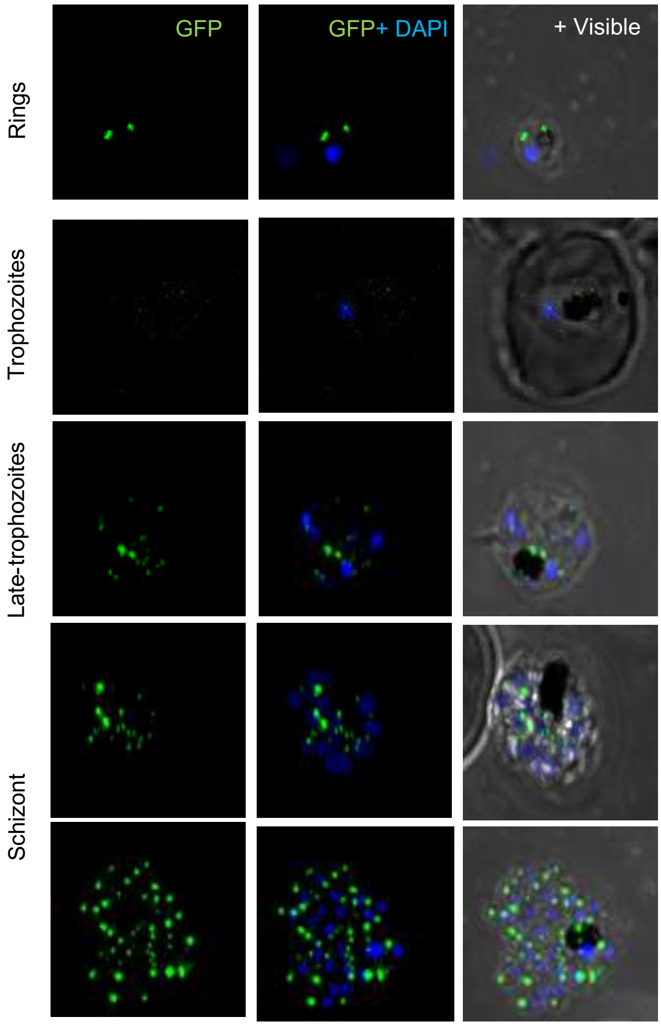


**Figure S2.** Confocal microscopy images of live transgenic parasites expressing *Pf*Dyn2-GFP fusion protein at different developmental stages during asexual blood stage cycle.

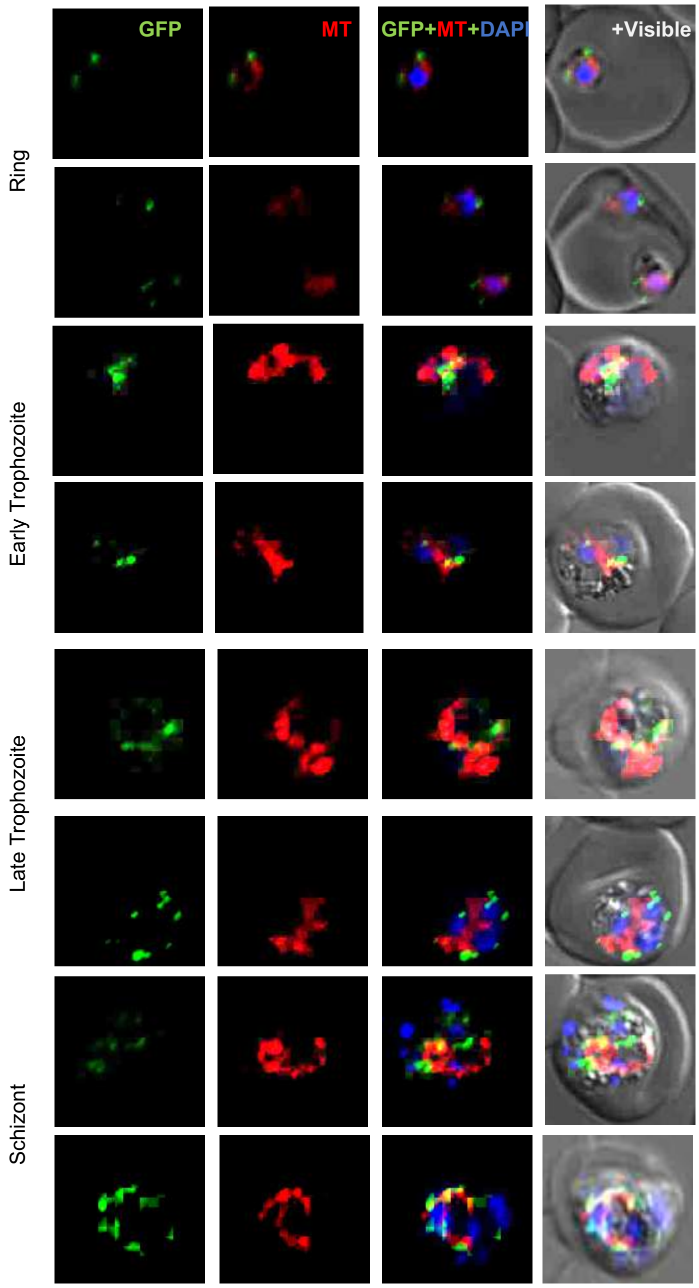


**Figure S3.** Stage-specific staining of parasite mitochondria to compare with localization of *Pf*DYN2. The fluorescence image of *Pf*DYN2-GFP parasites was stained with MitoTracker (red). Parasites were stained with DAPI and the parasite was visualized by a confocal microscope.

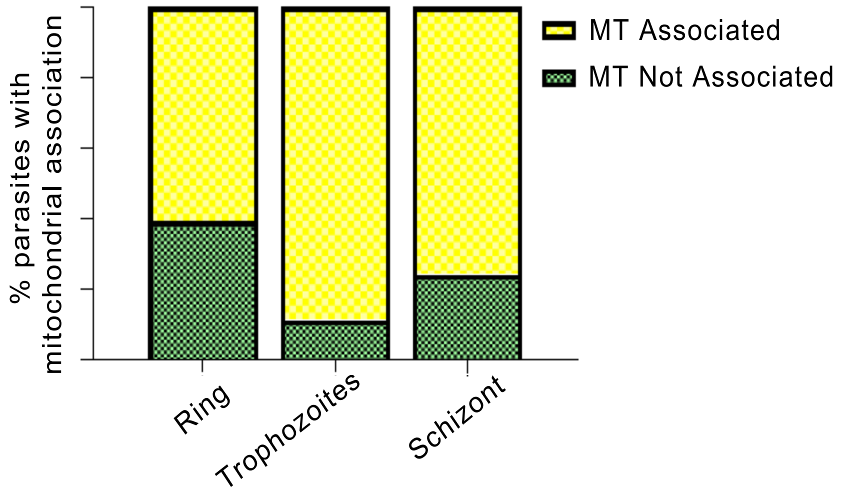


**Figure S4.** *Pf*Dyn2-GFP tagged fusion protein associated with Mitochondria stained with Mito-Tracker (red) were counted at various stages (ring, trophozoite, and schizont $n$=50 each).

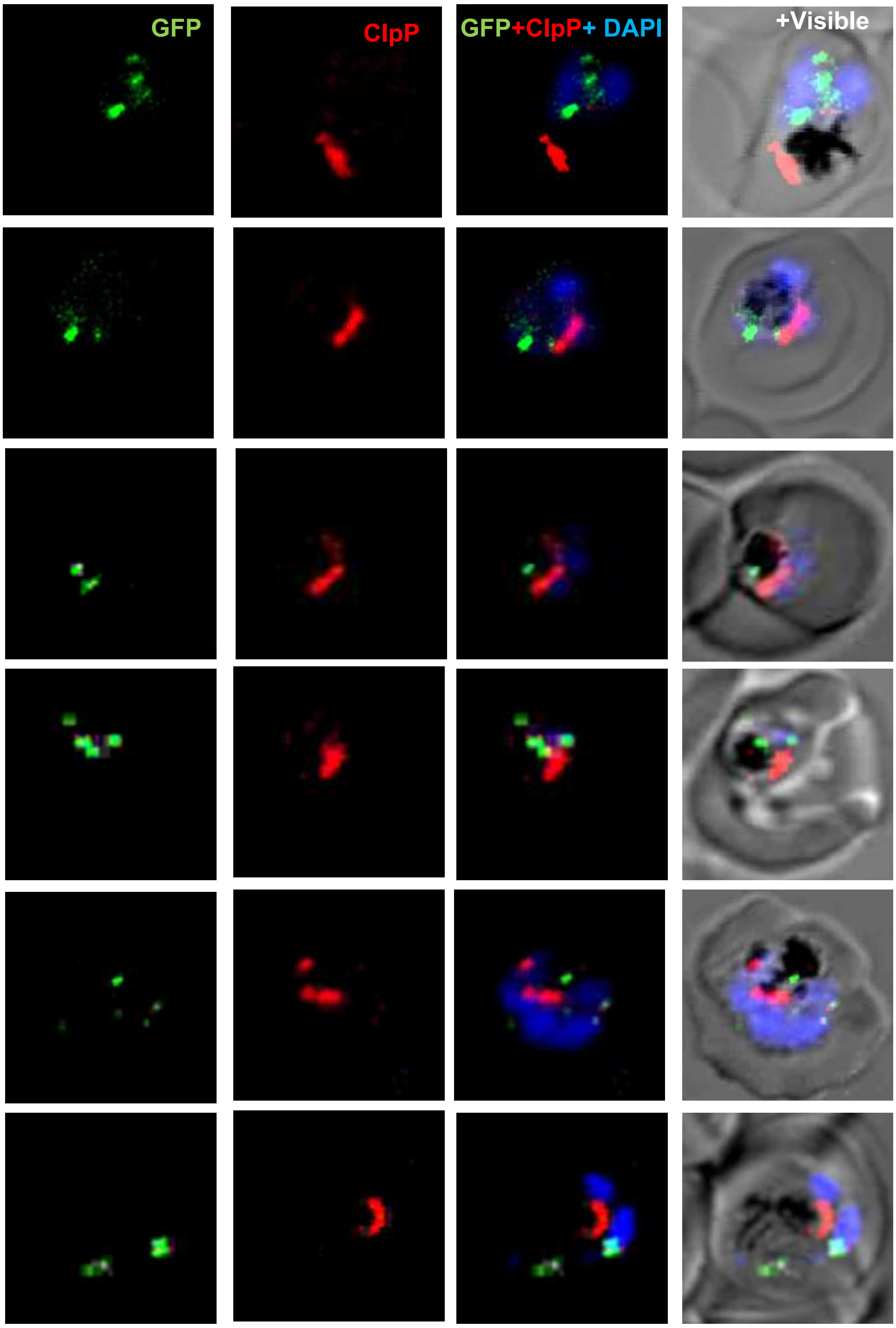


**Figure S5.** Labelling of parasite organelles, apicoplast to compare with localization of *Pf*Dyn2. Fluorescence images of transgenic parasites immuno-stained anti-*Pf*ClpP antibody. The PfDyn2-GFP punctate structure is seen as distinct from organelles labelling. Pearson's coefficient if less than 0.2 in both cases, which is not significant.

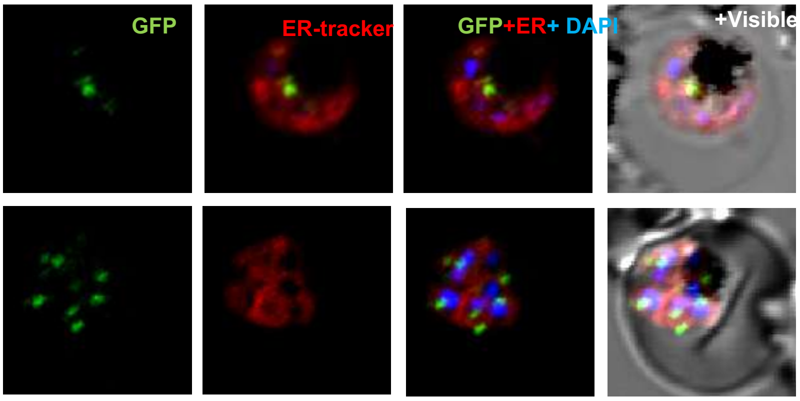


**Figure S6.** ***Pf*****Dyn2-GFP fusion protein does not associates with ER in the parasite**. Fluorescence images of trophozoites stage transgenic parasites expressing *Pf*Dyn2-GFP stained with ER Tracker (red). Pearson's coefficient if < 0.2.

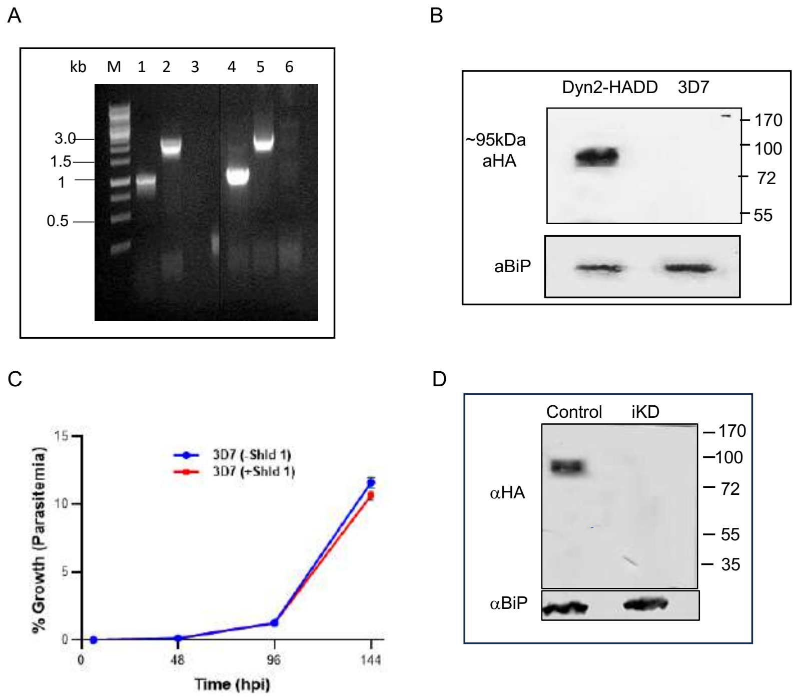


**Figure S7.** (A) PCR -based analyses to confirm integration of plasmid in the target gene locus using total DNAs of the transgenic parasite culture (purified clonal parasite population) and wild type 3D7 parasite lines, locations of primers are marked in the schematic: lane 1 and 4 (primers 1078A and 1234A) show amplification in both transgenic parasites with integrated plasmid and the wild-type parasites; lane 2 and 5 (primers 1466A and 1236A) showing amplification only in the parasites with integration; lane 3 and 6 (primers 1222A and 1223A) show amplification in both integrant or wild type parasites. (B) Western blot analysis of lysate of transgenic and wild-type parasites using anti-HA antibody. The fusion protein band (~95kDa) was detected in the transgenic parasites, PfDyn2-HADD line (lane 1) and not in the 3D7 parent parasite line (lane 2). Blot ran in parallel with an equal amount of the same sample, probed with anti-BiP antibody was used as a loading control. (C) Synchronous wild-type (3D7) parasites at the ring stage incubated with Shield-1 ligand and calculated the parasite percentage growth for three consecutive cycles. Graph showing Shld -1 is not toxic for the parasites. (D) Western blot analysis showing the reduction in the fusion protein and 84hpi, PfDyn2-HA, in transgenic parasites grown with Shld or without Shld1 drug (control and iKD).

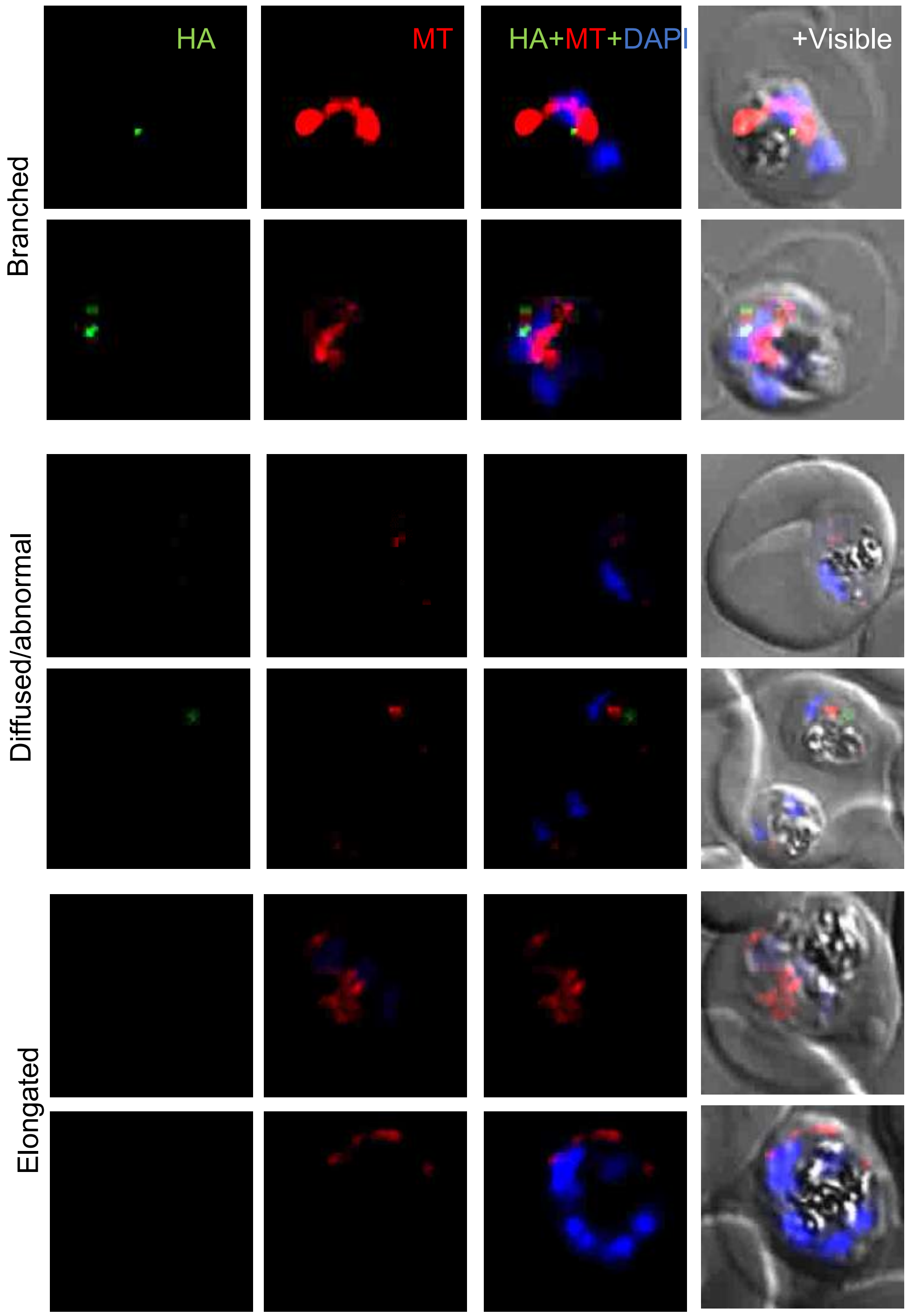


**Figure S8. Effect of downregulation of *Pf*Dyn2 at 27-36 hpi levels on growth and segregation of parasite mitochondrion.** Fluorescent microscopic images of transgenic *Pf*Dyn2-HA-DD parasites grown in the absence of Shld-1 drug *Pf*Dyn2-iKD sets, immuno-stained with anti-HA (green) antibodies and co-stained with MitoTracker dye (red).

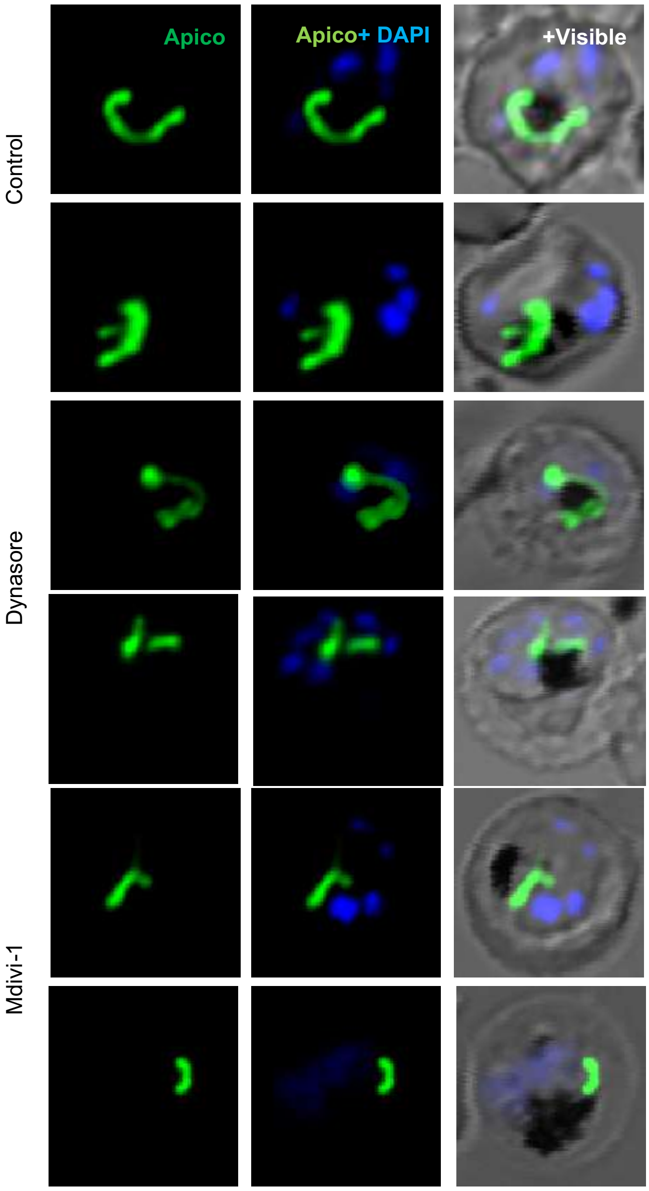


**Figure. S9. Effect of dynamin-specific inhibitors on Apicoplast D10-ACP GFP labeled transgenic parasite lines.** GFP labeled ACP-D10 apicoplast parasite lines treated at young trophozoite stages (27 hpi) with dynasore GTPase inhibitor (40 µM), Mdivi-1( 50 µM) or solvent (control) for 3 h. Fluorescent microscopic images of parasites from drug-treated and control sets were stained with DAPI (blue) and parasites were analyzed by confocal microscopy.

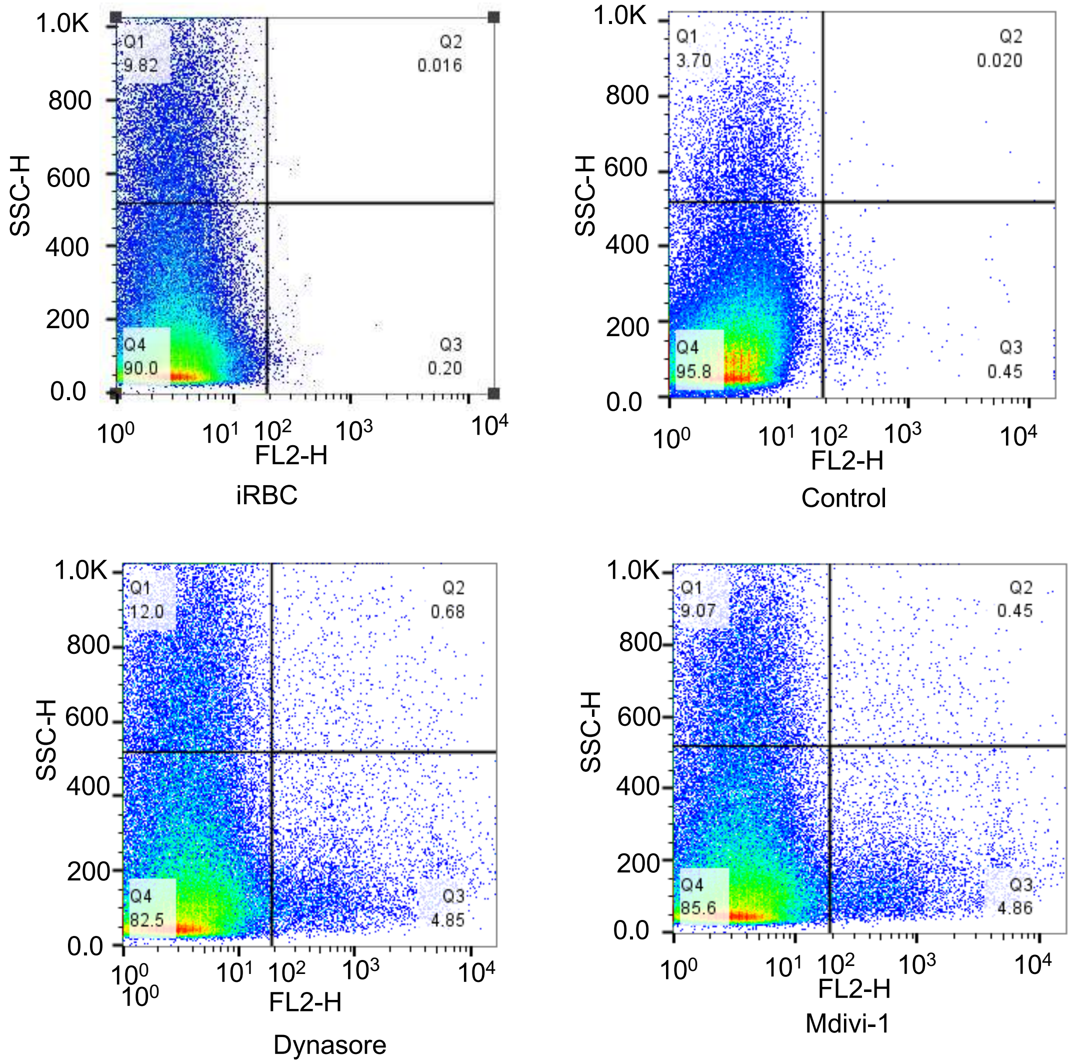


**Figure S10. Dynamin-specific inhibitors block mitochondrial segregation and induce Mitochondrial ROS production:** Dot plot showing significant induction of MioSOX red labeling (FL-2) in parasites after PfDyn2-iKD as compared to control parasites.
